# Targeted CRISPRi screening reveals unexpected resilience across the RNA polymerase III transcriptome

**DOI:** 10.64898/2026.09.24.754042

**Authors:** Boris Simeonov, Zeynep Veryeri, Yvette Faas, Alan Gerber

**Affiliations:** Department of Neurosurgery, Amsterdam UMC, Vrije Universiteit Amsterdam, Amsterdam, The Netherlands; Cancer Center Amsterdam, Cancer Biology and Immunology, Amsterdam UMC, Amsterdam, The Netherlands

**Keywords:** RNA polymerase III (Pol III), tRNA, CRISPR interference (CRISPRi), glioblastoma, non-coding RNA, functional screening

## Abstract

Increased RNA polymerase III (Pol III) activity and tRNA abundance are widely linked to cancer cell growth, yet the functional requirement for individual Pol III genes and core components remains unclear, in part due to the difficulty of achieving gene-specific perturbation of highly conserved loci. Here, we developed an inducible CRISPR interference platform and a custom single-guide RNA (sgRNA) library enabling gene-specific targeting of Pol III-transcribed genes and Pol III machinery. Genome-wide screening identified several Pol III dependencies in diploid fibroblasts and HEK293T cells, including multiple initiator methionine tRNA genes among the strongest fitness dependencies. Unexpectedly, glioblastoma models remained largely insensitive to repression of both individual Pol III genes and core Pol III components, despite efficient target repression. These findings establish a general strategy for gene-specific interrogation of conserved Pol III genes and indicate that glioblastoma models tolerate extensive perturbation of Pol III genes and machinery.

**Highlights:**

- A genome-wide CRISPRi library enables functional interrogation of individual Pol III genes.
- Contribution to the mature tRNA pool does not predict functional requirement.
- Individual tRNA genes exhibit context-dependent effects and opposing effects on cellular fitness.
- Glioblastoma models tolerate extensive perturbation of Pol III genes and machinery.

## Introduction

Pol III transcribes a diverse repertoire of small non-coding RNAs, including transfer RNAs and other structured RNAs that are essential for protein synthesis and gene regulation. Because these molecules support essential biosynthetic processes, Pol III genes have traditionally been regarded as housekeeping genes^1^. This view is consistent with their relatively compact promoter architecture compared to Pol II genes. Pol III genes are organized into three promoter types, with type I and type II promoters located within the transcribed region and type III promoters positioned upstream of the transcription start site. Transcription initiation relies on a defined set of factors, including TFIIIA, TFIIIC, and TFIIIB, which assemble at these elements to recruit Pol III, defining discrete regulatory regions flanking or overlapping the transcription unit. However, increasing evidence indicates that dysregulation of Pol III outputs, particularly mature tRNA pools, can reshape translational programs and contribute to disease development, such as cancer and neurological disorders^2,3^. For instance, specific tRNAs, such as tRNA^Glu(TTC)^ and tRNA^Arg(CCG)^, have been identified as metastasis promoters in breast cancer by enhancing the stability and ribosome occupancy of transcripts enriched for their cognate codons^4^. Moreover, a loss of a single brain-specific *TRR-TCT4-1* gene increases seizure susceptibility in mice, demonstrating that disruption of a single Pol III gene can contribute to disease phenotypes^5^. Similarly, copy number variations induced by targeting individual tRNA^Phe(GAA)^ isodecoder genes in mice have been shown to directly impair mammalian embryonic development and disrupt balanced, tissue-specific translation^6^. These observations demonstrate that perturbations of individual genes can influence cellular processes. However, it remains unclear whether such effects reflect a small number of functionally limiting loci or whether Pol III output is largely buffered by the collective activity of highly redundant tRNA pools. Resolving this distinction is essential for understanding how Pol III contributes to proliferation and disease^7–9^.

Distinguishing gene-specific functions from redundant Pol III output has remained challenging. Many Pol III genes, especially tRNAs, belong to multi-copy gene families with nearly identical sequences, complicating locus-specific targeting with standard methods. In humans, over 600 tRNA genes are categorized into various isotypes, many sharing full sequence identity, thus making it challenging to target individual genes without causing off-target effects^10,11^. Functional redundancy extends beyond genomic sequence similarity, as multiple tRNAs can decode the same codon through wobble base pairing. Consequently, perturbation of a single locus may be buffered by related gene copies, overlapping decoding capacity, and the plasticity of tRNA decoding.

Pooled CRISPR-Cas methodologies allow for the systematic classification of human genes into distinct functional categories and biological processes^12^. However, non-coding RNAs can be challenging to functionally disrupt via CRISPR-Cas9-mediated mutagenesis^13^. In the context of Pol III genes, these approaches are further constrained by the short length and extensive sequence similarity of many loci. CRISPR-Cas9-mediated disruption of anticodon sequences has been used to perturb tRNA function, but this approach operates at the isodecoder level and does not enable gene-specific targeting of individual loci^14^. RNA interference (RNAi) faces similar limitations, as identical or highly similar sequences across tRNA or other Pol III gene copies hinder locus-specific perturbation. The stable folding and the extensive nucleobase modifications of Pol III transcripts further limit the effectiveness of RNAi ^15^. CRISPR interference (CRISPRi) provides an alternative strategy for perturbing non-coding genes by repressing transcription without inducing DNA breaks. Although CRISPRi has been successfully applied to individual tRNA loci^16^, its extension to genome-wide studies of the Pol III transcriptome has been limited by the lack of validated targeting strategies capable of achieving efficient and locus-specific repression across the diverse classes of Pol III genes.

To systematically assess the contribution of individual Pol III genes and core Pol III components to cellular fitness, we first established generalizable CRISPRi targeting principles for efficient gene-specific repression of highly conserved Pol III loci. We then developed an inducible CRISPRi platform and a custom sgRNA library targeting Pol III–transcribed genes, as well as Pol III transcription factors and subunits. Using pooled dropout screens across non-cancer and two model glioblastoma cell lines, we systematically assessed the impact of Pol III perturbation on cell fitness. Repression of Pol III genes and core transcriptional components impaired fitness in non-cancer contexts, with multiple individual initiator methionine tRNA (iMet) genes emerging as the most impactful targets. In contrast, glioblastoma cells exhibited minimal sensitivity, as neither individual Pol III genes nor the core transcriptional components robustly impacted cell fitness despite comparable or greater knockdown efficiency. These findings reveal a pronounced, context-dependent resilience to both gene-specific and global Pol III perturbation.

## Results

### Developing a CRISPRi targeting strategy for Pol III genes

Previous work has proposed that mammalian cells maintain a broadly expressed housekeeping tRNA pool that supports core translational demands^7,14,17^ suggesting that a subset of highly expressed tRNAs provides core translational capacity and may thus disproportionately contribute to cellular fitness. Whether this relationship extends to individual genomic loci remains unknown. Individual tRNA genes contribute unequally to mature anticodon pools, and these contributions can vary across cell types^7^. For loci with uniquely assignable mature sequences (**Figure S1A**), this allows individual gene contributions to be quantified and raises the question of whether the dominant contributors to a mature tRNA pool are also the most important for cellular fitness. Addressing this question requires efficient repression of individual Pol III loci. However, the high sequence similarity among tRNA genes complicates locus-specific perturbation using conventional CRISPR-Cas9 approaches, as sgRNAs targeting gene bodies frequently match multiple loci^18^.

To enable gene-specific repression of Pol III loci, we established an inducible CRISPRi system using stable cell lines expressing doxycycline-inducible dCas9-KRAB-MeCP2, in which sgRNA-directed dCas9 recruits transcriptional repression domains to defined genomic loci without inducing DNA cleavage^19^ (**Figure S1B**). We selected this enhanced CRISPRi system because of its high repression efficiency in HEK293T cells, allowing us to target both Pol III-transcribed loci and Pol II-transcribed genes encoding components of the Pol III machinery. To guide sgRNA design, we first examined genome-wide binding profiles of Pol III transcription factors in ENCODE datasets relative to gene body-scaled annotated tRNA genes. ChIP-seq peak density profiles, representing the distribution of peak centers relative to the annotated start of mature tRNAs, showed that TFIIIC subunits peaked around the 5′ boundary and were distributed across the transcribed region, consistent with binding to internal promoter elements. In contrast, TFIIIB-associated peaks were predominantly enriched within ∼60 bp (**Figure 1A, upper panel**) of the 5’ boundary. These profiles define discrete regulatory regions flanking and overlapping Pol III transcription units.

**Figure 1.**
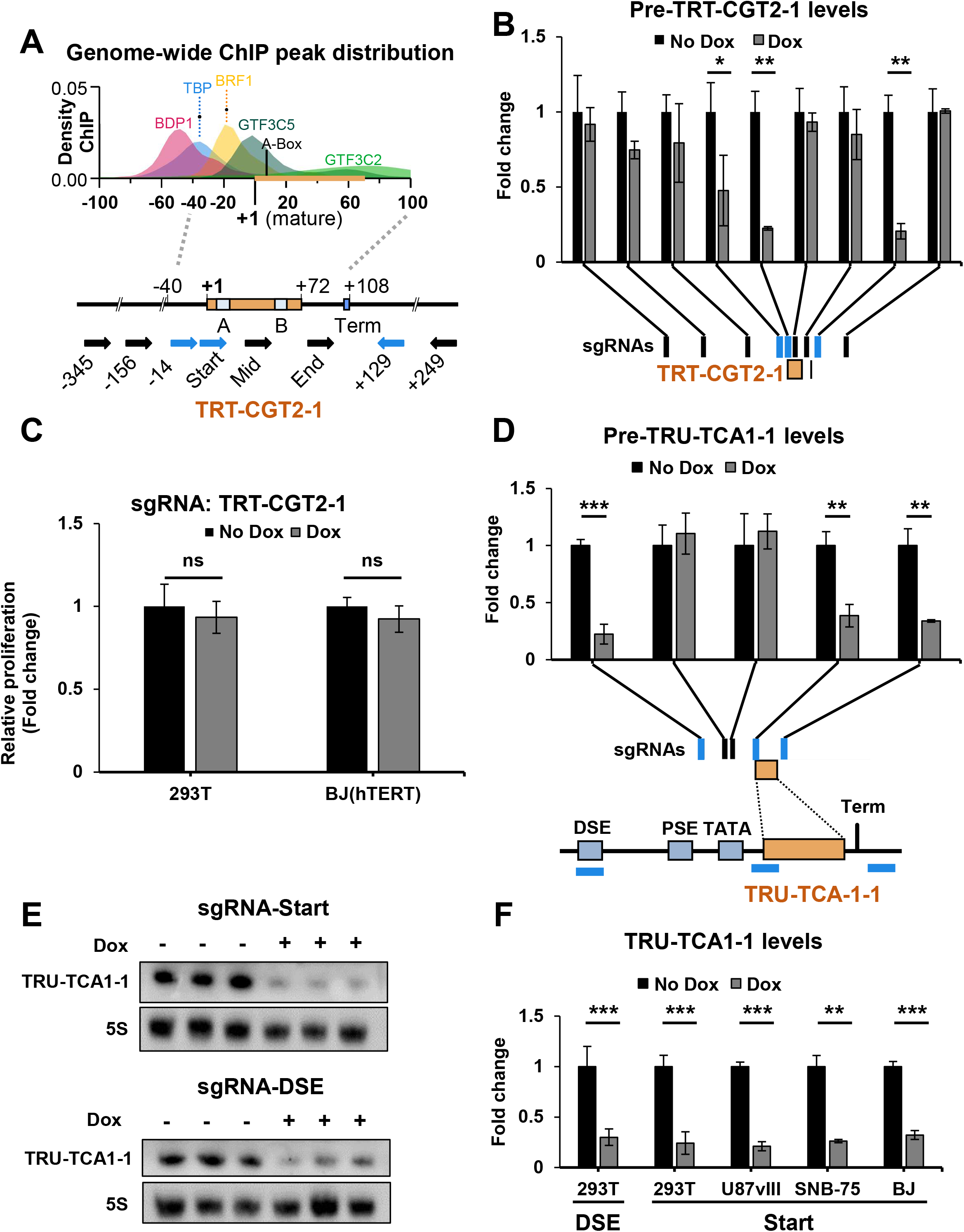
Experimental definition of CRISPRi targeting regions for Pol III genes. **A**) Genome-wide distribution of Pol III transcription-factor ChIP-seq peaks relative to the annotated 5′ boundary of mature tRNA genes (upper panel), and schematic representation of the *TRT-CGT2-1* locus showing the positions of sgRNAs used for tiling (lower panel). **B**) RT-qPCR analysis of pre-TRT-CGT2-1 levels following 48 h of doxycycline-induced CRISPRi using sgRNAs positioned across the locus. Transcript levels were normalized to PGK1 and expressed relative to the corresponding untreated condition. The lower schematic indicates sgRNA positions, with effective guides highlighted in blue. Data are shown as mean ± SD (n=3). Statistical significance was assessed using Welch’s t-test; *p < 0.05, **p < 0.01. **C**) Cell proliferation following repression of *TRT-CGT2-1* in cells expressing a 5′-proximal (“Start”) sgRNA, the dominant contributor to the mature tRNAThr(CGT) pool in HEK293T cells. Data shows average ± SD. AlamarBlue fluorescence was expressed relative to untreated cells cultured in the presence of Dox for 7 days in HEK293T (n = 6) and BJ-hTERT (n = 8) cells. **D**) Same as B) but with sgRNAs targeting the *TRU-TCA1-1* locus. **E**) Northern blot analysis of mature TRU-TCA1-1 following 7 days of CRISPRi using sgRNAs targeting the 5′-proximal (“Start”) or DSE regions (n=3). 5S rRNA was used as a reference RNA. **F**) Quantification of TRU-TCA1-1 Northern blots from HEK293T, U87vIII, SNB-75, and BJ-hTERT cells following 7 days of CRISPRi targeting the indicated regulatory region (Figure 1E **and Figure S1D**; n = 3). Values were normalized to 5S rRNA and expressed relative to untreated cells. Data are shown as mean ± SD; **p < 0.01, ***p < 0.001 (Welch’s t-test).

Based on this organization, we systematically evaluated sgRNA placement across Pol III loci. We therefore selected *TRT-CGT2-1* for method development because it represents the dominant contributor to the mature tRNA^Thr(CGT)^ pool (**Figure S1A**), allowing simultaneous optimization of CRISPRi targeting and assessment of whether dominant contributors to mature tRNA pools indeed represent functional dependencies. We first tiled nine sgRNAs across upstream, gene body, and downstream regions of the type II Pol III gene *TRT-CGT2-1* (**Figure 1A, lower panel**). Repression efficiency varied strongly with sgRNA position. The strongest effects were observed for guides targeting the 5′-proximal region, including the A box, and for guides positioned immediately downstream of the transcription terminator. In contrast, sgRNAs positioned within most of the gene body, including upstream of the terminator, or at upstream and downstream positions >100 bps, showed little or no activity. The absence of detectable repression at distal sites argues against substantial long-range spreading of KRAB-MeCP2-mediated repression under these conditions and instead indicates that effective repression is confined to regions proximal to Pol III regulatory elements (**Figure 1B**). Repression was maintained over extended doxycycline induction, with sustained reduction of both precursor and mature tRNA levels observed after up to 9 days of treatment (**Figure S1C**), indicating stable CRISPRi-mediated repression over time.

Given that *TRT-CGT2-1* contributes more than 50% to the tRNA^Thr(CGT)^ pool in 293T cells (**Figure S1A**), we next assessed whether repression of this dominant locus affects cell growth. Despite efficient repression of precursor transcripts, no measurable effect on proliferation was observed in 293T and BJ-hTERT (**Figure 1C**). Thus, contribution to the mature tRNA pool alone was not predictive of functional importance, motivating a systematic genome-wide assessment of Pol III gene dependencies.

To further evaluate if the observed sgRNA positional effects generalize across Pol III promoter architectures, we applied a similar targeting to the type III Pol III gene *TRU-TCA1-1*, including targeting of its upstream regulatory region, the distal sequence element (DSE) ^20,21^. As observed for *TRT-CGT2-1*, strong repression was achieved by sgRNAs targeting the 5′-proximal and terminator-associated regions, while targeting the DSE was similarly effective (**Figure 1D**). In contrast, sgRNAs positioned within the short intervening region between the DSE and the 5’-proximal regions showed little to no activity. These observations indicate that efficient CRISPRi repression is highly position-dependent and can be achieved by targeting distinct regulatory regions. Importantly, the lack of repression from intervening positions argues against extensive spreading of KRAB-MeCP2-mediated repression and instead suggests that effective CRISPRi requires targeting at or near specific regulatory elements.

Reduction of mature tRNA levels following *TRU-TCA1-1* targeting was confirmed by Northern blot analysis (**Figure 1E and 1F**), indicating that CRISPRi-mediated repression of precursor transcripts is reflected at the level of mature RNA. These effects were reproducible across multiple cell lines, including fibroblasts and glioblastoma models, with consistent repression observed for sgRNAs targeting 5′-proximal and upstream regulatory regions (**Figure 1F and S1D**).

Finally, to assess locus specificity, we targeted a Pol III-transcribed mammalian-wide interspersed repeat (MIR) gene located in the first intron of the POLR3E gene^22,23^, ∼800 bp from the TSS. CRISPRi targeting reduced MIR transcript levels without affecting POLR3E expression (**Figure S1E**), indicating that repression was restricted to the targeted Pol III locus without detectable interference with nearby Pol II transcription.

Together, these results identify 5′-proximal and terminator-associated regions as robust targeting windows for CRISPRi-mediated repression of Pol III loci. Across distinct promoter architectures and genomic contexts, repression remained restricted to defined regulatory regions and was sufficient to reduce both precursor and mature RNA levels. These observations provide a generalizable strategy for efficient and locus-specific targeting of Pol III genes.

### sgRNA library design for genome-wide targeting of Pol III genes

Having established positional rules for efficient repression of Pol III genes across distinct promoter architectures, we next asked whether these principles could be generalized to construct a genome-wide CRISPRi library for systematic interrogation of the Pol III transcriptome.

sgRNA selection was guided by the positional constraints defined in **Figure 1**. For each locus, sgRNA targeting windows were defined relative to the annotated gene coordinates. A 5′-proximal window (−40 to +20 relative to the annotated 5′ end) was directly defined from gene annotations. Because transcription termination sites are not annotated, we next systematically identified terminator sequences across the Pol III transcriptome by sequence analysis to enable gene-specific targeting in this second active window. Most loci contained canonical poly(T) terminators located approximately 10–20 bp downstream of the mature tRNA sequence (**Figure S2B**). These annotations enabled definition of a terminator-associated targeting window extending from −45 to +15 relative to the predicted termination site (**Figure 2A**). These windows correspond to the regions of maximal CRISPRi activity identified in **Figure 1**. sgRNAs were selected based on predicted specificity and sequence constraints, excluding guides with low specificity scores or homopolymeric stretches.

**Figure 2.**
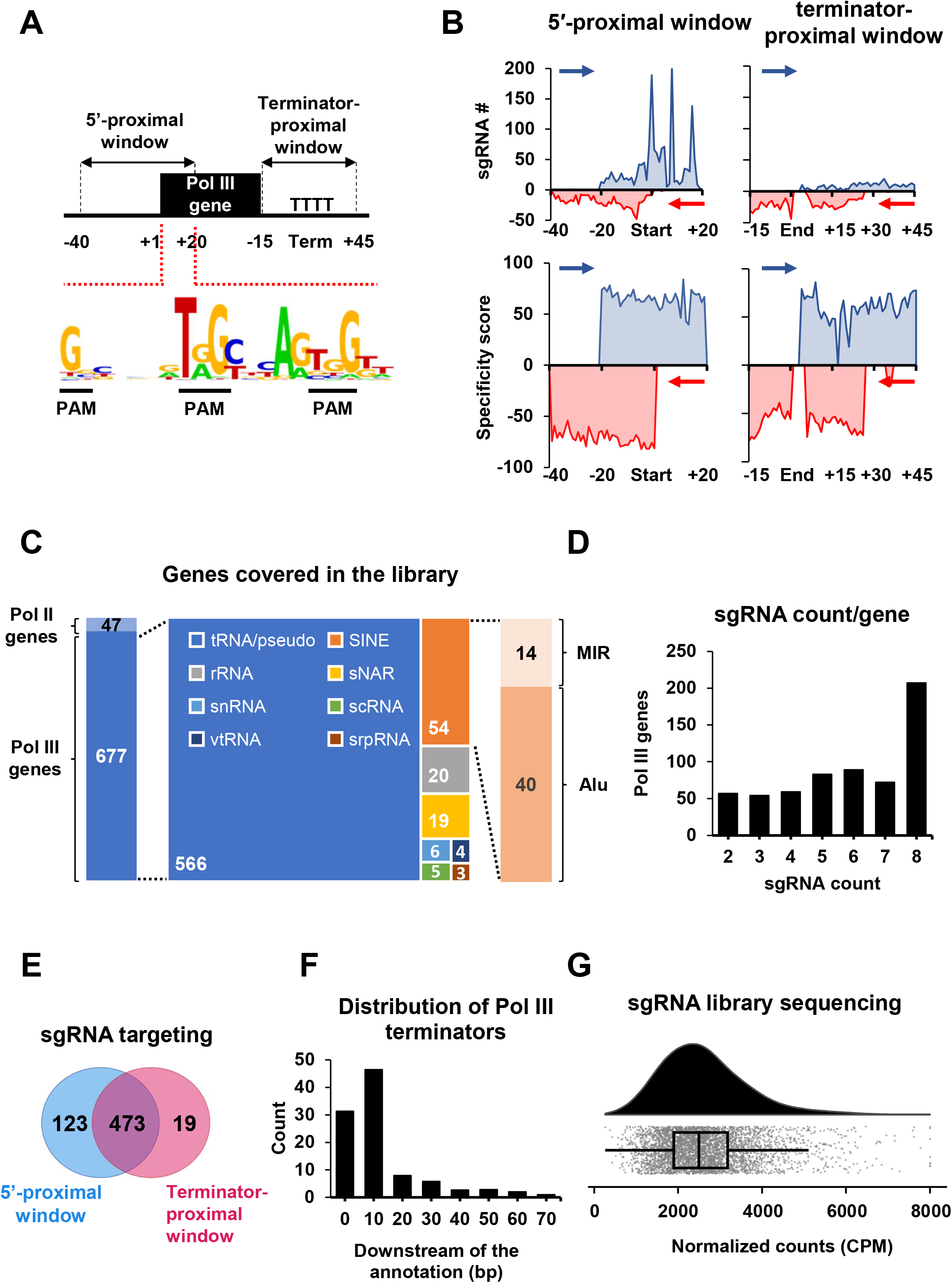
Design and composition of the genome-wide Pol III CRISPRi sgRNA library. **A**) Schematic of the sgRNA design strategy showing the 5′-proximal and terminator-proximal targeting windows defined from the tiling experiments in Figure 1. Recurrent PAM positions within the 5′ region, which includes the conserved A-Box in type II promoters, are indicated below the schematic. **B**) Distribution of sgRNA density and specificity scores across the 5′-proximal and terminator-proximal targeting windows. Blue and red traces indicate the end position of sgRNAs in the forward and reverse orientations, respectively. **C**) Composition of the final sgRNA library, showing the number of targeted Pol III-transcribed loci, Pol III machinery factors, and positive-control genes. Pol III-transcribed loci are further grouped by RNA class. **D**) Distribution of the number of sgRNAs assigned per targeted Pol III locus. **E**) Overlap of Pol III loci targeted in the 5′-proximal and terminator-proximal windows. **F**) Distribution of predicted Pol III terminator positions relative to the annotated 3′ boundary of the mature transcript. Terminators were identified by sequence analysis and are plotted by distance downstream of the annotated gene boundary. **G**) Distribution of normalized sgRNA read counts in the plasmid library following next-generation sequencing. Counts are shown as CPM and truncated at 8,000 CPM for visualization. The embedded box plot indicates the median and interquartile range.

Because many Pol III genes share identical mature sequences or extensive sequence homology, loci that could not be uniquely targeted were either excluded or intentionally grouped for shared perturbation. Grouped sgRNAs were classified into those targeting gene families (single sgRNA targeting multiple loci) and those with additional off-target matches (**Figure S2A**), enabling both gene-specific and multi-locus perturbation within the Pol III transcriptome.

The targeting windows identified in **Figure 1** were compatible with dense sgRNA coverage across the Pol III transcriptome. Within the 5′-proximal window, sgRNAs were strongly enriched at discrete positions corresponding to recurrent protospacer adjacent motifs (PAMs), including two conserved PAM-containing sites within the A box and a frequent PAM occurrence near the annotated 5′ end of the gene, thereby placing sgRNAs directly within regions associated with the highest CRISPRi activity. In contrast, distribution across the terminator-associated window was more uniform, excluding regions overlapping terminator sequences (**Figure 2A and 2B**). sgRNAs selected within both targeting windows exhibited comparable specificity score distributions, indicating that positional enrichment was not driven by relaxed selection criteria (**Figure 2B and S2C**). The final library comprised a total of 4110 sgRNAs targeting 566 tRNA genes, 54 SINE elements, and 57 additional Pol III–transcribed non-coding RNAs, covering the majority of uniquely targetable and active Pol III loci, as well as 36 Pol III subunits and transcriptional regulators (**Figure 2C; Table S2**). Positive control sgRNAs targeting essential genes and validated CRISPRi-sensitive loci were included alongside 249 non-targeting controls. Most genes were targeted by up to eight sgRNAs, with a minimum of two sgRNAs per gene to enable robust gene-level inference (**Figure 2D**). A subset of 473 loci was targeted in both 5′-proximal and terminator-associated windows, enabling direct comparison of positional effects within individual genes (**Figure 2E**). The majority of the terminators were located 10 bps downstream of the gene annotation (**Figure 2F**). Following cloning and lentiviral library generation, sequencing of the plasmid pool confirmed uniform representation of sgRNAs without major skewing (**Figure 2G**). The resulting library combines broad coverage of the Pol III transcriptome with locus-specific targeting and provides a platform for genome-wide functional screening of Pol III genes and associated factors.

### Functional screening identifies Pol III dependencies not predicted by tRNA abundance

To benchmark the genome-wide Pol III CRISPRi library and establish screening conditions, we first performed a pooled dropout screen in HEK293T cells. Following transduction and selection, cells were split into doxycycline-treated (D) and untreated (ND) populations, with an initial reference sample collected at time point zero (T0). Cells were cultured for approximately 20 population doublings prior to genomic DNA extraction and deep sequencing of the integrated sgRNA cassettes to quantify sgRNA abundance (**Figure 3A**). This strategy was chosen to allow depletion of stable Pol III transcripts, including mature tRNAs with long half-lives^24^ thereby allowing detection of fitness effects caused by sustained transcriptional repression.

**Figure 3.**
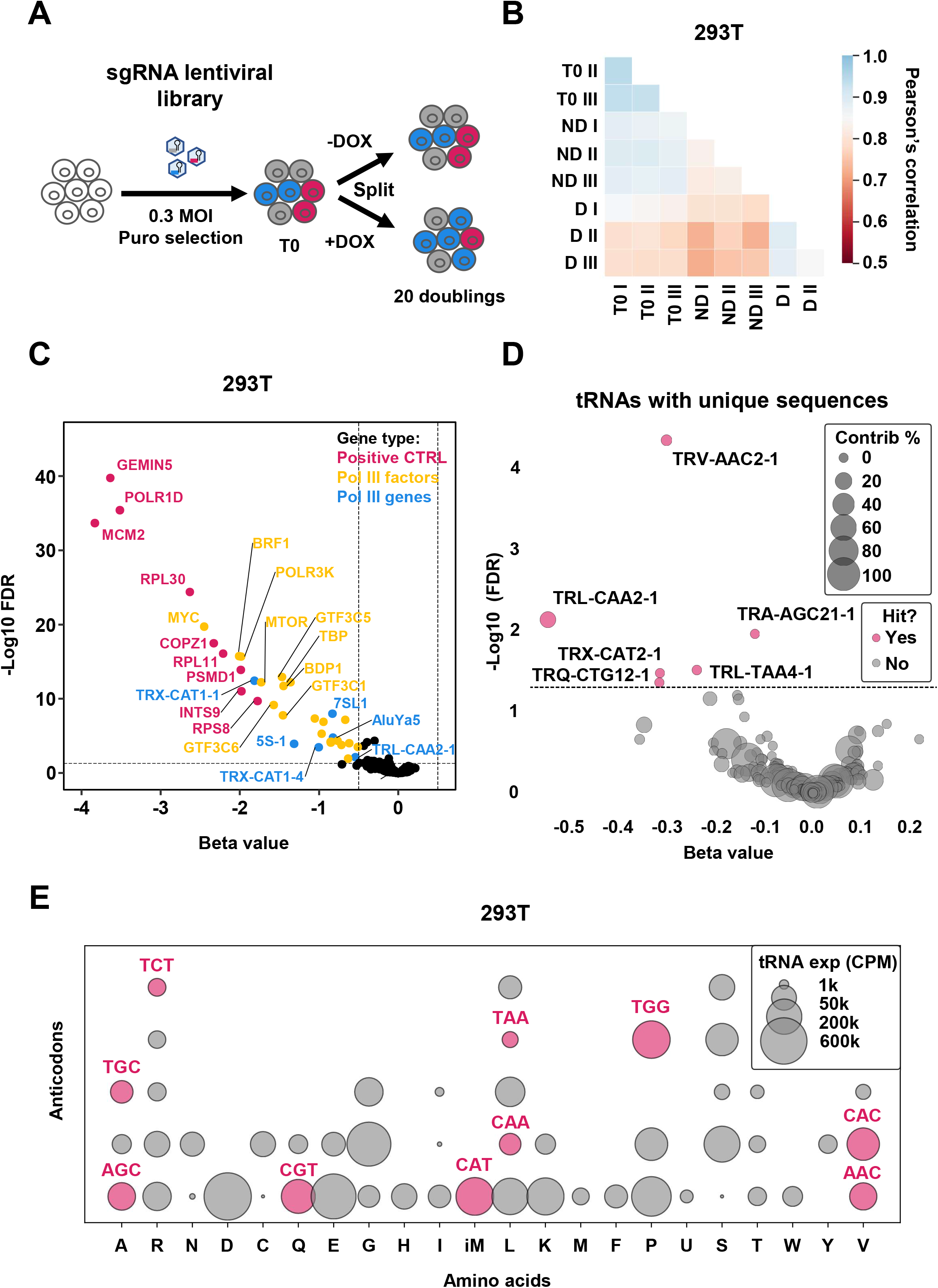
Genome-wide Pol III CRISPRi screening identifies dependencies not predicted by tRNA abundance. **A**) Experimental workflow for pooled CRISPRi screening of the Pol III sgRNA library. Following transduction and selection, a T0 reference sample was collected, and cells were cultured with or without doxycycline for approximately 20 population doublings before sgRNA abundance was determined by sequencing. **B**) Pairwise Pearson correlation of sgRNA abundance across biological replicates and experimental conditions in the HEK293T screen. D, doxycycline-treated; ND, untreated; T0, initial reference population. **C**) Volcano plot showing gene-level effects in the HEK293T screen (n = 3). Beta scores indicate estimated fitness effects, with negative values representing depletion and positive values enrichment following CRISPRi induction. The horizontal dashed line indicates FDR = 0.05. Vertical dashed lines indicate β = ±0.5. **D**) Screening results for tRNA loci with uniquely assignable mature sequences. Bubble size indicates the contribution of each locus to its corresponding mature anticodon pool. The horizontal dashed line indicates FDR = 0.05. **E**) Relative abundance of mature tRNA anticodon pools grouped by amino acid. Bubble size represents anticodon abundance (CPM); color indicates whether the corresponding anticodon family contained at least one significant tRNA gene hit in the HEK293T screen (red, ≥1 hit; grey, no hit).

A pilot screen performed with HEK293T cells revealed that biological replicates showed high reproducibility across all conditions, with strong correlation between replicates and clear separation of dox-treated samples from T0 and untreated (ND) controls (**Figure 3B; Table S3**). Independent analyses using either T0 or ND as a reference yielded highly concordant results (**Figure S3A**) and highly similar sgRNA abundance distributions across samples (**Figure S3B**). Based on this concordance, subsequent screens were analyzed using T0 as the primary reference.

Screen performance was further supported by the consistent depletion of positive control sgRNAs in the 293T cell line (**Figure 3C; Table S4**), indicating robust CRISPRi activity and reliable detection of fitness effects. In addition, a family of 17 5S rRNA co-targeted genes was identified as strongly depleted, consistent with their essential role in ribosome function and with efficient targeting of Pol III genes.

Among individually targeted tRNA genes, several initiator methionine tRNA (tRNA^iMet^, TRX) loci emerged as prominent depleted hits, including *TRX-CAT1-1*, *TRX-CAT1-4* and *TRX-CAT2-1*. Because tRNA^iMet^ is uniquely required for translation initiation, sensitivity to perturbation of this tRNA family may be expected. Nevertheless, the repeated identification of independent tRNA^iMet^ loci contrasts with the limited representation of most other tRNA families among significant hits, with tRNA^iMet^ genes accounting for three of the thirteen significantly depleted tRNA genes (FDR < 0.05). Additional non-tRNA depleted loci were also detected, indicating that the screen captures functional requirements beyond positive controls and highly redundant multicopy Pol III genes. The lack of a proliferative phenotype following repression of *TRT-CGT2-1*, despite its contribution of more than half of the mature tRNA^Thr(CGT)^ pool (**Figure 1C and S1A**), suggested that abundance alone may not predict functional importance. We therefore asked whether this observation generalized across the Pol III transcriptome. Because individual gene contributions can only be resolved for tRNAs with unique mature sequences, this analysis was restricted to uniquely assignable loci. Comparison of screening outcomes with mature-pool contribution revealed no clear relationship between abundance and fitness impact (**Figure 3D**). None of the significantly depleted tRNA genes represented dominant contributors to their respective anticodon pools, whereas several of the largest contributors showed no detectable fitness effect. Thus, genes that account for a substantial fraction of mature tRNA abundance are not necessarily functionally limiting for proliferation.

To determine whether this observation extended beyond individual genes, we examined the relationship between anticodon abundance and screening outcome across the tRNA transcriptome. Anticodons containing significant hits were distributed across a broad range of expression levels and were not preferentially associated with either highly abundant or weakly expressed tRNA pools (**Figure 3E**). Together, these results indicate that neither gene-level contribution to mature tRNA pools nor overall anticodon abundance is sufficient to predict functional dependency, highlighting the value of functional screening for identifying limiting components of the Pol III transcriptome.

Together with the *TRT-CGT2-1* analysis, these results demonstrate that functional dependencies within the Pol III transcriptome cannot be inferred from tRNA abundance measurements alone, establishing the need for direct functional interrogation across diverse cellular contexts.

### Glioblastoma models exhibit reduced sensitivity to perturbation of individual Pol III genes

We next applied genome-wide CRISPRi screening across additional non-cancer (BJ-hTERT, **Table S5**) and SNB-75 as well as U87vIII glioblastoma (**Table S6** and **S7**, respectively) cell lines to determine how Pol III gene dependencies vary across cellular contexts. This design allowed us to distinguish common Pol III dependencies from cell type-specific requirements. Consistent depletion of positive-control sgRNAs was observed in all four screens, demonstrating robust screen performance across cellular contexts (**Figure 4A**). In addition, multiple Pol III factors, including Pol III subunits and transcriptional regulators, were identified among the strongest depleted hits in BJ-hTERT and HEK293T cells, whereas surprisingly few showed comparable depletion in the glioblastoma models. All four screens showed a strong bias toward depleted rather than enriched hits, indicating that repression of Pol III-associated genes more frequently impaired than enhanced cellular fitness (**Tables S4-S7**).

**Figure 4.**
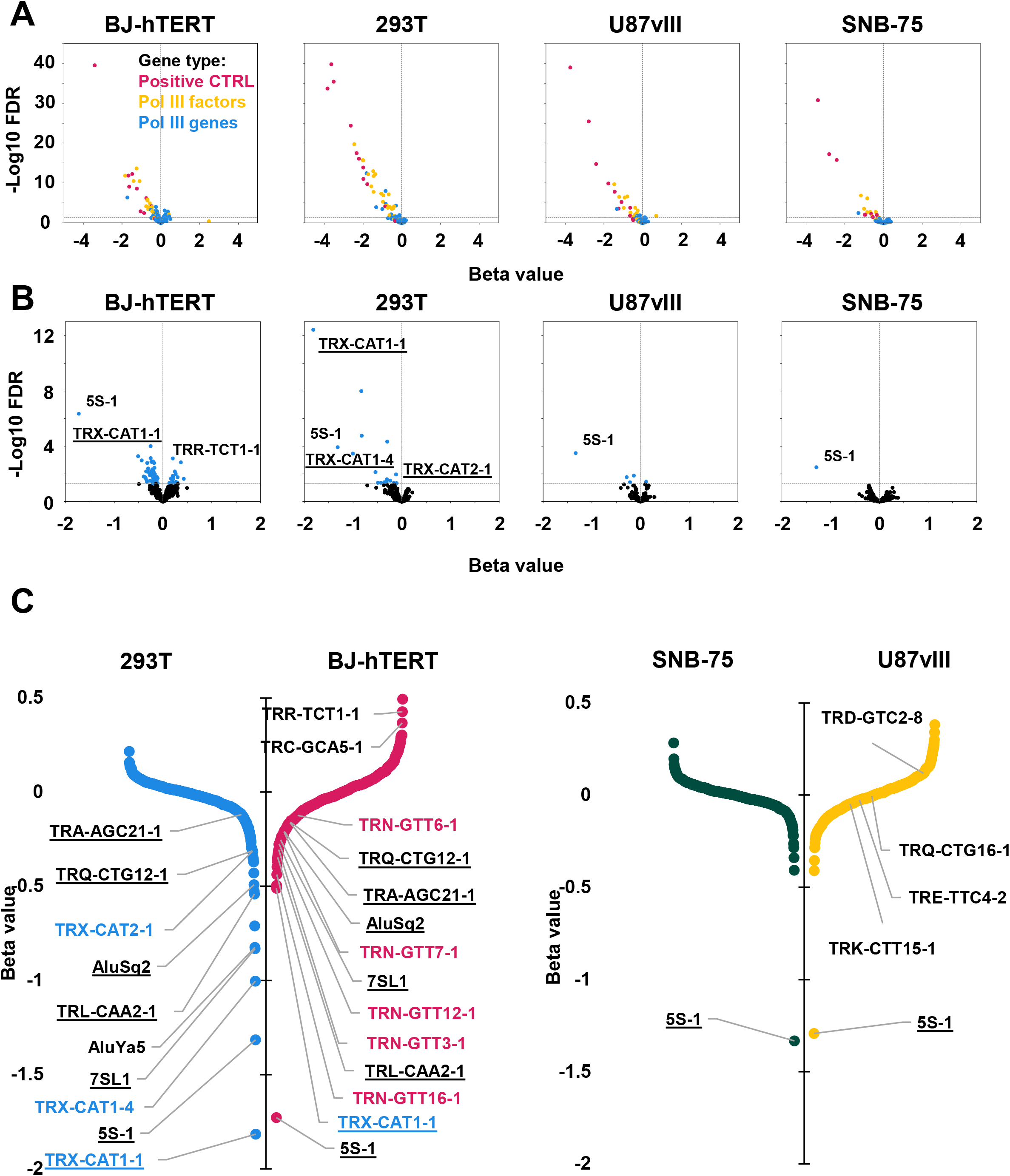
Glioblastoma models exhibit markedly reduced dependency on individual Pol III genes. **A**) Gene-level screening results across BJ-hTERT, HEK293T, U87vIII, and SNB-75 cells (n = 3). Pol III-transcribed genes are shown in blue, Pol III machinery factors in yellow, and positive-control genes in red. The horizontal dashed line indicates FDR = 0.05. **B**) Screening results restricted to Pol III-transcribed genes after exclusion of positive controls and Pol III machinery components. Selected hits are labeled, including *TRX-CAT* loci (underlined), the 5S rRNA family, and *TRR-TCT1-1*. The horizontal dashed line indicates FDR = 0.05. **C**) Rank plots of beta scores for all screened Pol III-transcribed genes in each cell line. Negative β scores indicate depletion and positive scores indicate enrichment following CRISPRi induction. Underlined labels indicate genes identified as significant hits in at least two cell lines. *TRX-CAT* loci are highlighted in blue and *TRN-GTT* loci in magenta. 5S-1 denotes the co-targeted 5S rRNA family.

To focus specifically on Pol III-transcribed genes, we reanalyzed screening results after excluding positive controls and Pol III machinery components (**Figure 4B**). This analysis revealed striking differences between cell types. BJ-hTERT fibroblasts displayed a broad set of significant hits with generally modest effect sizes, whereas HEK293T cells exhibited a smaller number of dependencies associated with larger fitness effects. In contrast, few Pol III genes reached significance in U87vIII cells and virtually none in SNB-75 cells. Quantification of significant Pol III gene hits further illustrated this trend (**Figure S4A**). BJ-hTERT fibroblasts exhibited the broadest dependency landscape, with 64 unique Pol III gene hits and an additional seven shared with HEK293T cells. HEK293T cells displayed only 10 unique hits, whereas U87vIII and SNB-75 exhibited just three and no unique dependencies, respectively. Thus, detectable Pol III gene dependencies were most numerous in diploid fibroblasts, fewer in transformed HEK293T cells, and nearly absent in glioblastoma models. Importantly, the shared depletion of 5S rRNA genes across all four screens demonstrates that CRISPRi-mediated repression of Pol III genes can generate measurable fitness defects in every cellular context examined. The marked reduction in significant hits observed in glioblastoma cells is therefore unlikely to reflect ineffective CRISPRi-mediated repression.

Examination of the strongest dependencies further highlighted distinct patterns across cell types (**Figure 4C**). *TRX-CAT1-1* emerged as the strongest depleted Pol III gene in both BJ-hTERT and HEK293T cells. By contrast, neither *TRX-CAT1-1* nor any other tRNA^iMet^ locus reached significance in U87vIII or SNB-75 cells, despite their rapid proliferative capacity. Fibroblasts exhibited a broader dependency profile spanning multiple tRNA families, including several tRNA^Asn(GTT)^, whereas HEK293T hits were dominated by a smaller number of high-effect dependencies. Additional non-tRNA Pol III loci were also identified, including *RN7SL1* and a shared Alu element in both non-cancer cell lines, indicating that detectable fitness effects extend beyond tRNA genes.

These findings demonstrate that dependence on individual Pol III genes is highly context-dependent. Whereas non-cancer cells displayed multiple shared and unique Pol III gene dependencies, glioblastoma cells remained remarkably insensitive to repression of individual Pol III genes despite robust screen performance.

The recurrent identification of *TRX-CAT1-1* as the strongest shared dependency in both non-cancer cell lines, together with the absence of a detectable fitness dependency in glioblastoma, prompted its further characterization.

### Individual tRNA gene perturbations produce context-dependent fitness effects

We first examined the behavior of individual sgRNAs targeting *TRX-CAT1-1*. Guides targeting both the 5′-proximal and terminator-associated regions were consistently depleted following doxycycline treatment, with stronger depletion of 5′-proximal than terminator-associated guides (**Figure S5A**).

To validate this result, selected sgRNAs were tested individually in 293T cells. Targeting either the 5’-proximal or the terminator-associated windows led to efficient repression at the pre-tRNA level within 48 hours (**Figure 5A**). At the mature tRNA level, 5′-proximal targeting reduced the total tRNA^iMet(CAT)^ pool by ∼50% (**Figure 5B**), demonstrating that *TRX-CAT1-1* contributes substantially to the initiator methionine tRNA pool. In contrast, the terminator-associated sgRNA resulted in a more modest change that did not reach statistical significance at the bulk level (**Figure 5B**), despite a pronounced reduction at the pre-tRNA levels (**Figure 5A**). Nevertheless, both targeting strategies produced significant fitness defects in the screen despite their differential effects on the mature tRNA^iMet^ pool.

**Figure 5.**
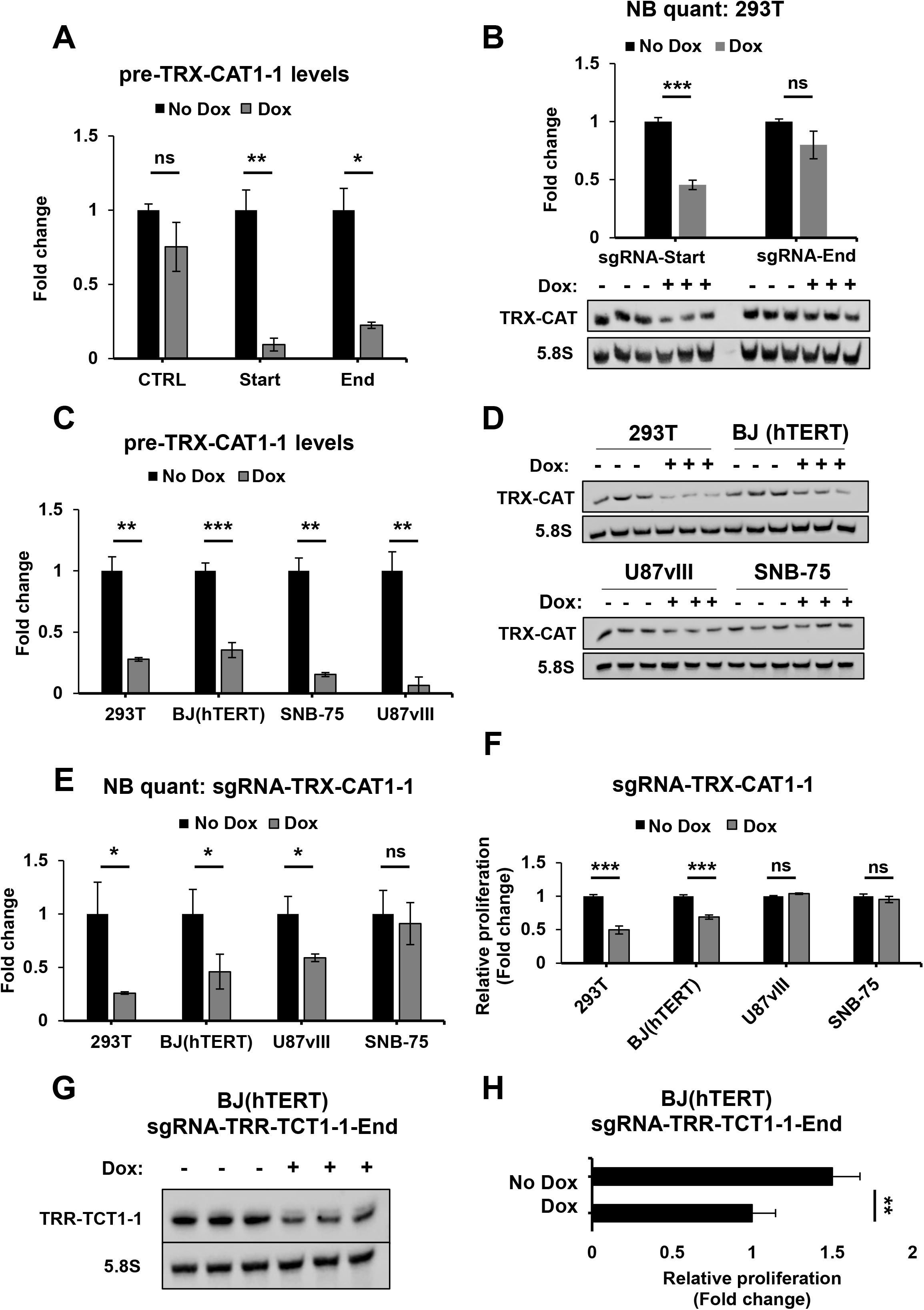
*TRX-CAT1-1* repression produces context-dependent effects on mature tRNA abundance and cell fitness. **A**) RT-qPCR validation of pre-TRX-CAT1-1 repression in HEK293T cells following 48 h of doxycycline-induced CRISPRi using sgRNAs targeting the 5′-proximal (“Start”) or terminator-associated (“End”) regions (n=3). Transcript levels were normalized to PPIA. **B**) Northern blot analysis of the total mature tRNA^iMet(CAT)^ pool following 7 days of *TRX-CAT1-1* repression using 5′-proximal or terminator-associated sgRNAs in HEK293T cells. 5.8S rRNA was used as a loading reference. Quantification of the Northern blot is shown above (n=3). **C**) RT-qPCR analysis of pre-TRX-CAT1-1 repression following 48 h of doxycycline induction in HEK293T, BJ-hTERT, SNB-75, and U87vIII cells using a 5′-proximal sgRNA (n=3). Transcript levels were normalized to PPIA. **D**) Northern blot analysis of mature tRNA^iMet(CAT)^ following 7 days of *TRX-CAT1-1* repression in the indicated cell lines. 5.8S rRNA was used as a loading reference. **E**) Quantification of the Northern blots shown in D (n=3), normalized to 5.8S rRNA. **F**) Relative proliferation following 7 days of *TRX-CAT1-1* repression in the indicated cell lines, assessed by alamarBlue (n=5). **G**) Northern blot analysis of mature TRR-TCT1-1 following 7 days of terminator-associated CRISPRi in BJ-hTERT cells. 5.8S rRNA was used as a loading reference. **H**) Relative proliferation of BJ-hTERT cells following 7 days of *TRR-TCT1-1* repression (n=5). For quantitative analyses, values are expressed relative to the corresponding untreated (no doxycycline) condition. Data represent average ± SD from independent biological replicates. Statistical significance was assessed using two-sided Welch’s t-test (ns, p > 0.05; *p < 0.05; **p < 0.01; ***p < 0.001).

Although *TRX-CAT1-1* pre-tRNA repression was comparable across all tested cell lines (**Figure 5C**), its impact on mature tRNA levels differed substantially. Non-cancer cell lines exhibited a pronounced reduction in the mature tRNA^iMet(CAT)^ pool, whereas glioblastoma cell lines showed reduced or minimal changes, with a partial decrease observed in U87vIII and little to no effect in SNB-75 cells (**Figure 5D and 5E**), suggesting that glioblastoma cells can buffer transcriptional repression at the level of the mature tRNA pool. Despite efficient precursor repression, *TRX-CAT1-1* targeting impaired cell growth in non-cancer cells but had no measurable impact in glioblastoma models (**Figure 5F**). Importantly, differences in mature tRNA abundance alone did not explain the differential fitness response. In U87vIII cells, *TRX-CAT1-1* repression reduced mature tRNA^iMet(CAT)^ to an extent comparable to that observed in BJ-hTERT cells, yet did not impair proliferation. Thus, glioblastoma resistance cannot be explained solely by maintenance of the mature tRNA pool but also reflects greater tolerance to its reduction. Because *TRX-CAT1-1* lies only ∼250 bp from the essential Pol II gene *ILF2* (**Figure S5B**), we asked whether its fitness phenotype could result from collateral repression of *ILF2*. Although partial reduction of *ILF2* expression was observed in 293T cells at the RNA and protein levels (**Figure S5C-S5E**), this effect was not detected in BJ-hTERT cells, where *TRX-CAT1-1* repression also reduced fitness. As controlled overexpression of mature functional tRNAs is technically challenging and can require multicopy genomic transgenes to achieve substantial increases in the mature tRNA pool^25^ we instead directly tested the potential contribution of ILF2. Ectopic expression of ILF2 during CRISPRi targeting in 293T cells rescued the growth defect caused by direct *ILF2* repression but not that caused by *TRX-CAT1-1* targeting (**Figure S5F-S5H**), demonstrating that the *TRX-CAT1-1* fitness phenotype is not attributable to collateral *ILF2* repression.

The screens also identified a small number of tRNA loci whose repression was associated with increased rather than decreased fitness. We therefore selected *TRR-TCT1-1*, an enriched hit in BJ-hTERT cells, for independent validation. Terminator-associated targeting efficiently reduced the corresponding tRNA (**Figure 5G**) and reproducibly increased proliferation (**Figure 5H**), validating the direction of the screening phenotype.

These experiments validate the diverse fitness effects identified by the screen and reveal distinct responses to tRNA gene perturbation across cellular contexts. *TRX-CAT1-1* repression reduced fitness in HEK293T and BJ-hTERT cells but was tolerated in glioblastoma cells despite comparable repression of the precursor transcript. In glioblastoma cells, this resistance was associated with either maintenance of the mature tRNA^iMet(CAT)^ pool or tolerance of its reduction, demonstrating that the effect of tRNA gene repression on mature tRNA abundance does not alone determine the resulting fitness phenotype. Conversely, repression of *TRR-TCT1-1* increased fibroblast proliferation, demonstrating that individual tRNA genes can exert distinct and even opposing effects on cellular fitness.

### Resistance to Pol III perturbation in glioblastoma extends to the transcriptional machinery

The tolerance of glioblastoma cells to repression of individual Pol III genes suggested that these cells may possess substantial buffering capacity within the Pol III system. Under this scenario, perturbation of the Pol III transcriptional machinery would likewise be expected to have limited consequences for cellular fitness. To test this possibility, we examined the effects of targeting core components of the Pol III transcription machinery in all cell models. Consistent with the results obtained for Pol III genes, targeting core Pol III subunits and associated general transcription factors showed strong depletion in BJ-hTERT and HEK293T cells, whereas glioblastoma cells were relatively less sensitive (**Figure 6A and 6B**). In contrast, this reduced sensitivity was not observed for transcriptional regulators that participate in Pol III transcription but also have broader roles in Pol II transcription and transcriptional regulation, including MYC, JUN, TBP, and SNAPC components (**Figure S6A**). This distinction suggests that the reduced dependency of glioblastoma cells is preferentially associated with factors dedicated to the Pol III transcriptional machinery rather than with transcriptional regulators more broadly. We next asked whether differential sensitivity could simply reflect differences in the abundance or chromatin association of Pol III machinery across cell lines. Fractionation of cytoplasmic+nucleoplasmic and chromatin-associated protein pools revealed no clear correlation between Pol III factor levels and sensitivity to perturbation. While 293T cells exhibited higher overall levels of several Pol III components and BJ-hTERT cells showed lower levels, glioblastoma cell lines showed intermediate abundance without a corresponding increase in dependency (**Figure 6C**). These observations suggest that absolute levels of Pol III machinery do not explain the reduced sensitivity observed in glioblastoma cells.

**Figure 6.**
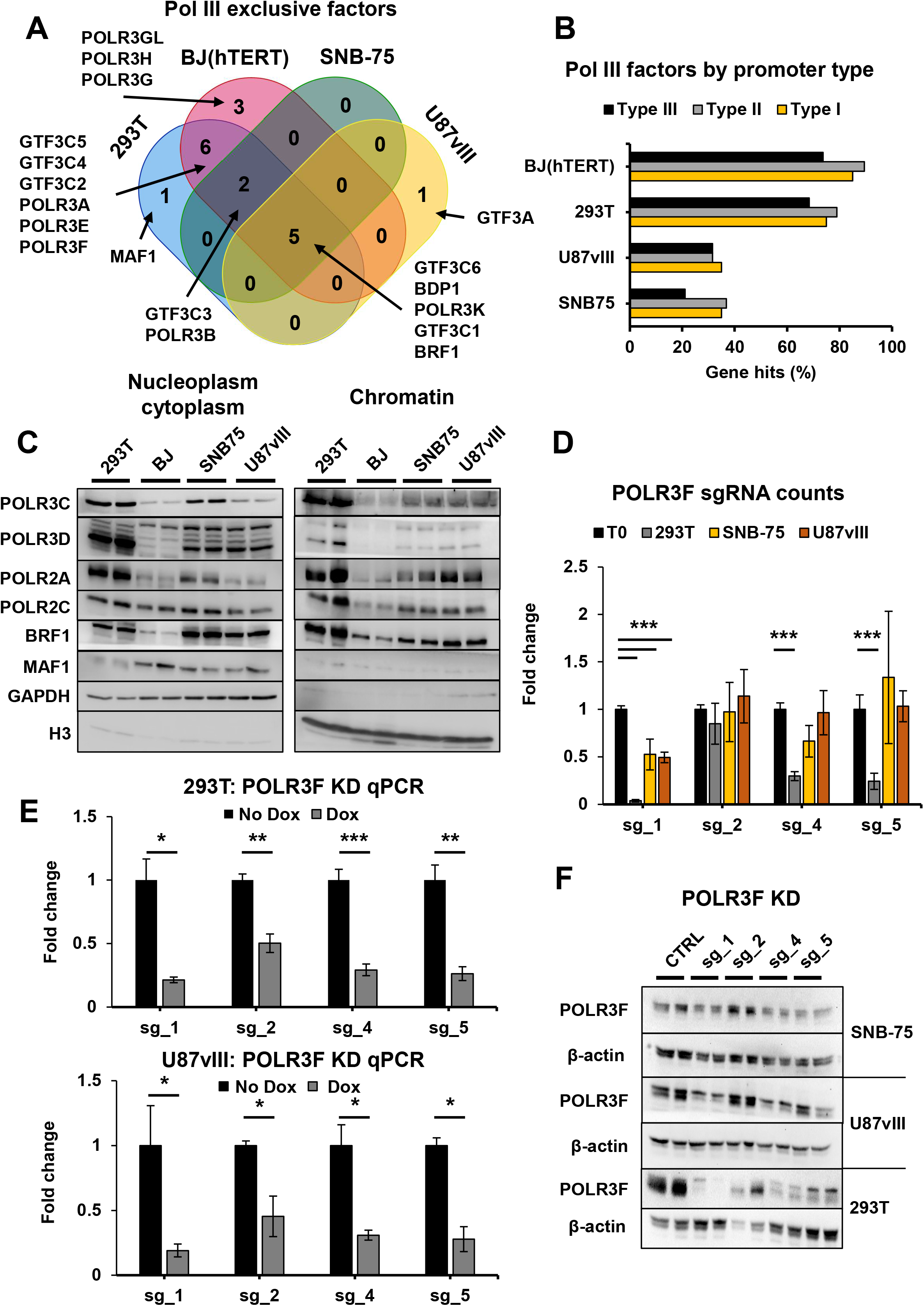
Reduced sensitivity to Pol III perturbation in glioblastoma extends to the transcriptional machinery. **A**) Venn diagram showing significant Pol III machinery dependencies identified across the four CRISPRi screens. **B**) Fraction of screened Pol III machinery factors identified as significant dependencies in each cell line, grouped according to the Pol III promoter types with which they are associated. **C**) Immunoblot analysis of Pol III machinery components in soluble cytoplasmic/nucleoplasmic and chromatin-associated fractions from the indicated cell lines. GAPDH and histone H3 were used as markers for the soluble and chromatin-associated fractions, respectively. **D**) Relative abundance of individual sgRNAs targeting POLR3F following CRISPRi screening in three cell lines (n=3). CPM-normalized sgRNA counts are expressed relative to the corresponding T0 population. **E**) RT-qPCR analysis of POLR3F repression following 7 days of doxycycline induction in HEK293T and U87vIII cells (n=3). POLR3F transcript levels were normalized to PGK1 and expressed relative to untreated cells. **F**) Immunoblot analysis of POLR3F protein following 7 days of CRISPRi using four independent POLR3F-targeting sgRNAs in HEK293T, U87vIII and SNB-75 cells (n=2). Doxycycline-treated cells lacking a POLR3F-targeting sgRNA were used as controls, and β-actin was used as a loading reference. For quantitative analyses, data represent mean ± SD from independent biological replicates. Statistical significance was assessed using two-sided Welch’s t-test (*p < 0.05; **p < 0.01; ***p < 0.001).

Likewise, puromycin incorporation revealed no clear relationship between global translation output and sensitivity to Pol III machinery perturbation (**Figure S6B**). In particular, 293T and U87vIII cells displayed comparable levels of puromycin incorporation after normalization, despite marked differences in sensitivity to Pol III perturbation, whereas SNB-75 cells exhibited lower overall incorporation without increased sensitivity. These observations indicate that global translation rates do not explain the differential response to Pol III perturbation across cell lines.

To directly validate the differential sensitivity of non-cancer and glioblastoma cells to perturbation of the Pol III machinery, we focused on POLR3F, a core Pol III subunit identified as a dependency only in non-cancer cells. Multiple sgRNAs targeting POLR3F showed differential depletion patterns across cell lines, with strong effects observed in non-cancer cells but limited impact in glioblastoma models (**Figure 6D**). Despite efficient reduction of POLR3F mRNA across all tested cell lines (**Figure 6E**), the extent of protein depletion varied between cell lines (**Figure 6F**), indicating that transcriptional repression was not uniformly propagated to the protein level. Importantly, independently of this buffering at the protein level, the substantial POLR3F protein depletion in U87vIII cells was not accompanied by a fitness defect (**Figure 6D**), indicating greater tolerance to perturbation in this cell line. This remarkably parallels the response to *TRX-CAT1-1* repression (**Figure 5**), where glioblastoma cells either buffered the effect of transcriptional repression on mature tRNA abundance or tolerated reduction of the mature tRNA pool. Together, these observations indicate that glioblastoma resilience to Pol III perturbation can occur both through maintenance of the targeted gene product and through tolerance of its depletion.

These results demonstrate that the reduced sensitivity of glioblastoma cells to Pol III perturbation extends beyond individual Pol III-transcribed genes to core components of the transcriptional machinery. This phenotype was not explained by differences in Pol III factor abundance or global protein synthesis and persisted despite efficient repression of POLR3F, indicating that glioblastoma cells tolerate substantial perturbation of the Pol III system without measurable fitness consequences.

## Discussion

### A genome-wide CRISPRi framework enables locus-resolved interrogation of the Pol III transcriptome

Earlier genome-wide CRISPR screens have largely focused on Pol II–transcribed genes, leaving the functional organization of the Pol III transcriptome largely unexplored.^19,26,27^. Previous studies have examined the consequences of broader Pol III perturbation, including through MAF1 perturbation^28^ or targeted tRNAs at the anticodon level^14^, but these approaches do not resolve the contributions of individual Pol III-transcribed loci.

Systematic functional analysis of Pol III genes has been limited by their high sequence similarity, which complicates locus-specific perturbation. For tRNAs, redundancy extends beyond genomic sequence similarity, as multiple gene copies can contribute to the same mature tRNA pool and distinct anticodons can provide overlapping decoding capacity through wobble interactions and modification-dependent decoding. Here, we established a genome-wide CRISPRi platform targeting Pol III genes and associated factors, enabling functional interrogation of the Pol III transcriptome at the locus level despite these constraints. By defining positional targeting rules, we show that efficient repression can be achieved from discrete regulatory regions, including the 5′-proximal region, the DSE of a type III promoter locus, and sequences immediately downstream of the transcription terminator, whereas intervening and most gene-body positions were largely ineffective. The sharp positional dependence of repression argues against extensive spreading of KRAB-MeCP2-mediated repression and instead indicates that efficient CRISPRi requires positioning at or near specific regulatory regions^29^.

Previous studies have established that KRAB-based CRISPRi can repress Pol III-transcribed genes^30^. Our positional analysis further suggests that the local organization of the Pol III transcription complex strongly influences CRISPRi efficacy. One possible explanation for the poor activity of most gene-body sgRNAs is that occupancy by TFIIIC and other components of the Pol III transcription machinery limits access or activity of the CRISPRi complex at these positions^31,32^. Efficient repression from promoter-proximal regulatory regions is readily compatible with interference with transcription complex assembly. More unexpected was the strong activity of sgRNAs positioned immediately downstream of the transcription terminator. One possible explanation is interference with facilitated recycling of Pol III^33,34^, which enables efficient re-initiation at highly transcribed tRNA genes. Disruption of this process could reduce repeated rounds of transcription without requiring direct targeting of promoter-proximal regions.

These findings establish practical targeting principles for CRISPRi of highly conserved Pol III-transcribed genes and enable their systematic functional interrogation at genome scale.

### Context-dependent resilience to perturbation of the Pol III system

Targeted validation of *TRX-CAT1-1* revealed that the relationship between transcriptional repression, mature tRNA levels, and cell fitness is strongly context-dependent. Despite comparable repression of the precursor transcript, the effect on the mature tRNA^iMet(CAT)^ pool differed substantially between cell lines. In SNB-75 cells, precursor repression produced little detectable change in mature tRNA abundance, indicating buffering between transcriptional repression and the mature RNA pool. Previous studies of MAF1 loss have shown that increased Pol III transcription does not necessarily result in proportional increases in mature tRNA abundance^35–37^, demonstrating that transcriptional output and steady-state mature tRNA levels can be uncoupled. Our findings reveal that such uncoupling can also occur following reduced transcriptional output, at least in SNB-75 cells. Importantly, however, maintenance of the mature tRNA pool was not required for resistance to *TRX-CAT1-1* repression in the second glioblastoma model. U87vIII cells instead tolerated a substantial reduction in mature tRNA^iMet(CAT)^ without measurable impairment of proliferation. Thus, the two glioblastoma models exhibited distinct responses to *TRX-CAT1-1* repression: maintenance of the mature tRNA pool in SNB-75 and tolerance of its reduction in U87vIII.

Models of translational regulation have proposed that tRNA abundance is coordinated with codon usage to optimize translation efficiency and support gene expression programs, particularly in proliferative and oncogenic contexts^35,38^. Consistent with this view, increased Pol III activity and elevated tRNA expression have been widely reported in cancer cells and are thought to support growth and biosynthetic demand^39–41^. However, our genome-wide analysis showed that tRNA abundance was a poor predictor of functional dependency. Neither the contribution of individual loci to mature anticodon pools nor overall anticodon abundance was associated with fitness effects. Notably, repression of *TRT-CGT2-1*, which contributes more than half of the mature tRNA^Thr(CGT)^ pool in HEK293T cells, did not measurably impair proliferation. Thus, quantitative contribution to the mature tRNA pool does not by itself determine whether an individual locus is functionally limiting. The same distinction between buffering and tolerance extended to the Pol III transcriptional machinery. Repression of core Pol III subunits and dedicated transcription factors produced strong fitness defects in BJ-hTERT and 293T cells but substantially weaker effects in glioblastoma models. For the Pol III subunit POLR3F, efficient mRNA repression was not uniformly propagated to the protein level, indicating buffering at the level of the final gene product. Importantly, substantial POLR3F protein depletion in U87vIII cells occurred without a corresponding fitness defect. This remarkably parallels the response to *TRX-CAT1-1,* suggesting that glioblastoma cells can either attenuate the effect of transcriptional repression on the final gene product or tolerate depletion once it occurs. The reduced sensitivity was not explained by differences in baseline Pol III factor abundance, chromatin association, or global translation rates, indicating that reduced sensitivity does not arise from increased transcriptional capacity or altered protein synthesis demands^42,43^. Moreover, the effect was preferentially associated with Pol III-dedicated machinery and was not observed for Pol III-associated regulators with broader transcriptional functions, arguing against a general resistance to perturbation of transcriptional regulators.

These observations reveal a striking uncoupling between perturbation of the Pol III system, abundance of its downstream products, and cellular fitness. Glioblastoma cells tolerated repression of individual Pol III genes, including initiator methionine tRNA loci, as well as perturbation of core Pol III machinery. Importantly, this resilience cannot be attributed to a single homeostatic mechanism. In some contexts, transcriptional repression was attenuated at the level of the mature RNA or protein product, whereas in others substantial depletion of the final product was tolerated without measurable consequences for proliferation. The molecular basis of this tolerance remains unclear and could involve excess functional capacity, altered turnover, compensatory regulation, or differences in cellular demand. Nevertheless, the ability to tolerate perturbation at multiple levels distinguishes the glioblastoma models from BJ-hTERT and 293T cells, in which comparable perturbations more readily produced fitness defects.

### Expression and functional dependency are uncoupled within the tRNA transcriptome

Previous studies have proposed that cancer cells rely on increased Pol III activity and selective upregulation of tRNAs to support proliferation and gene expression programs^44–46^. While such changes in tRNA abundance are well documented, our results show that increased abundance does not necessarily imply increased functional dependency. In the glioblastoma models examined here, individual tRNA genes and components of the Pol III machinery produced remarkably few fitness dependencies despite the rapid proliferation of these cells. More broadly, across the cell models tested, neither the contribution of individual tRNA loci to their mature anticodon pools nor overall anticodon abundance predicted their fitness effects.

Importantly, the absence of a simple relationship with abundance does not imply that individual tRNA genes are functionally equivalent. The screens identified discrete and strongly context-dependent dependencies. *TRX-CAT1-1* was the strongest shared Pol III gene dependency in BJ-hTERT and 293T cells, whereas several tRNA^Asn(GTT)^ loci were selectively required in fibroblasts. Conversely, repression of *TRR-TCT1-1* increased proliferation in BJ-hTERT cells, demonstrating that perturbation of an individual tRNA locus can exert fitness effects in either direction. Thus, gene-specific tRNA dependencies exist, but they cannot be inferred simply from the quantitative contribution of a locus to the mature tRNA pool.

The identification of multiple initiator methionine tRNA genes as dependencies is consistent with the specialized requirement for tRNA^iMet^ in translation initiation. However, even this dependency was context-dependent, with strong depletion of *TRX-CAT1-1* in BJ-hTERT and 293T cells but not in either glioblastoma model. Thus, the unique translational function of a tRNA family does not necessarily translate into a general dependency on its individual genomic loci. Conversely, despite contributing more than half of the mature tRNA^Thr(CGT)^ pool in 293T cells, repression of *TRT-CGT2-1* produced no measurable proliferation defect. Several loci identified as dependencies, in turn, were not dominant contributors to their respective mature pools. Functional importance therefore cannot be inferred simply by ranking tRNA genes according to expression or pool contribution. Taken together, our findings reveal multiple levels of uncoupling between tRNA expression and functional dependency. Basal abundance or contribution to the mature tRNA pool did not predict which loci were required for fitness, while repression of individual loci did not uniformly translate into depletion of the mature RNA. When mature tRNA levels were reduced, comparable changes could produce markedly different fitness outcomes across cellular contexts. These observations demonstrate that expression, perturbational response, and functional requirement represent distinct properties of individual tRNA loci and underscore the value of direct functional interrogation of the Pol III transcriptome at the single-locus resolution.

### Limitations of the study

Several limitations should be considered. First, while CRISPRi enables locus-specific repression, it does not provide precise control over the magnitude of tRNA depletion, limiting quantitative assessment of the thresholds at which individual Pol III genes may become functionally limiting. Consequently, loci that appear dispensable following partial transcriptional repression could still become limiting upon complete genetic loss. Orthogonal genetic deletion approaches will therefore be required to determine whether the observed dependencies extend to complete loss of individual loci. In addition, redundancy among tRNA genes and overlapping decoding capacity may mask dependencies that would emerge only upon simultaneous perturbation of multiple loci. Combinatorial targeting will therefore be required to systematically interrogate redundancy within tRNA pools. Second, although CRISPRi targeting was designed to maximize locus specificity, collateral effects on nearby Pol II-transcribed genes cannot be excluded for all loci. In the case of *TRX-CAT1-1*, partial repression of the adjacent *ILF2* gene was observed in 293T but not in other cell lines, and functional rescue experiments indicated that this did not account for the observed phenotype. Importantly, similar genomic configurations appear to be rare among positive hits, suggesting that co-targeting of protein-coding genes is unlikely to represent a general confounding factor. Nevertheless, potential effects on neighboring non-coding genes should therefore be considered when interpreting individual dependencies. Third, the 5′-proximal targeting window was defined relative to the annotated 5′ end of the mature tRNA rather than experimentally mapped transcription initiation sites, which are not available for most human tRNA loci. Variation in the length of 5′ leader sequences may therefore affect the position of individual sgRNAs relative to the transcriptional machinery and contribute to differences in targeting efficiency. Nevertheless, the enrichment of Pol III transcription-factor occupancy around the annotated 5′ boundary (**Figure 1A**), together with the experimentally defined positional activity of sgRNAs, supports this region as a practical targeting window.

Finally, the biological scope of the study is limited to proliferative fitness in four established cell lines under standard culture conditions. In particular, the reduced sensitivity to Pol III perturbation observed in U87vIII and SNB-75 should not be interpreted as a general property of glioblastoma. Additional patient-derived models will be required to establish the generality of this phenotype. Moreover, Pol III genes that are dispensable for proliferation under the conditions tested may become functionally limiting during differentiation, metabolic or nutrient stress, or other cellular states. Despite these limitations, the platform provides a foundation for systematic investigation of Pol III gene function across diverse biological contexts. Future applications could combine perturbation of related tRNA loci to resolve functional redundancy, define quantitative thresholds of tRNA availability, and extend functional screening to additional cellular states and disease-relevant models.

## Supplemental information

### Supplementary figure legends

**Figure S1.**
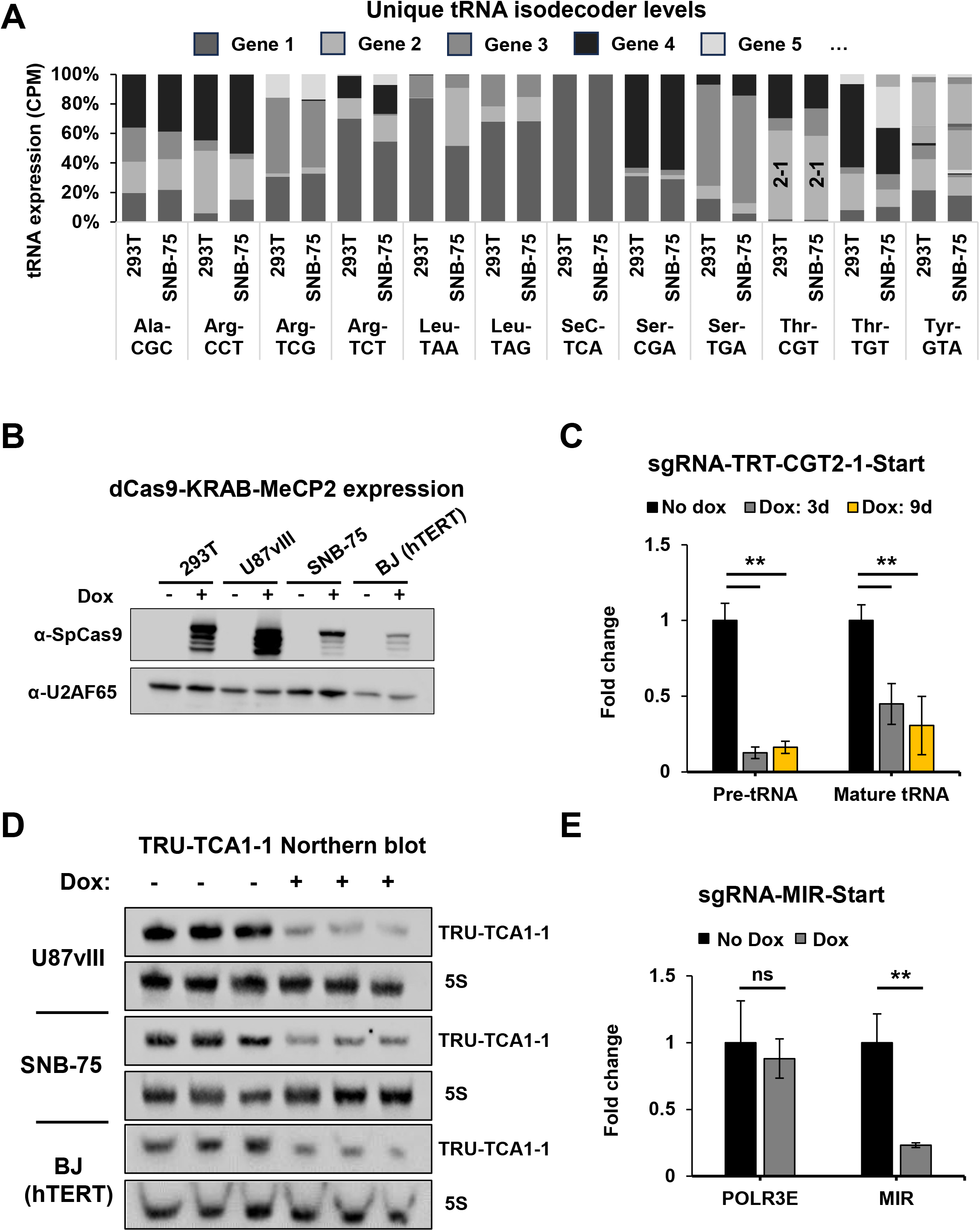
Related to Figure 1, **A**) Relative abundance of tRNA loci with uniquely assignable mature sequences and their contribution to the corresponding mature anticodon pool in HEK293T and SNB-75 cells. tRNA-seq data are from Scheepbouwer et al^17^. **B**) Western blot of the induction of dCas9-KRAB-MeCP2 expression after 24 h of doxycycline (dox) treatment in the cell lines used for the CRISPRi screen. U2AF65, loading control. **C**) Time course of *TRT-CGT2-1* repression, with pre-tRNA and mature tRNA quantified by RT-qPCR relative to *PGK1* mRNA in HEK293T cells, showing sustained knockdown. Data represent mean ± SD (n = 3). **D**) Northern blot of *TRU-TCA1-1* tRNA after one week of dox treatment. 5S rRNA, loading control (n = 3). **E**) RT-qPCR analysis of a Pol III-transcribed MIR element located within the first intron of *POLR3E* and of POLR3E mRNA following MIR-targeted CRISPRi. Transcript levels were normalized to PGK1 mRNA levels. For quantitative analyses, data represent mean ± SD from independent biological replicates. Statistical significance was assessed using two-sided Welch’s t-test (*p < 0.05; **p < 0.01; ***p < 0.001).

**Figure S2.**
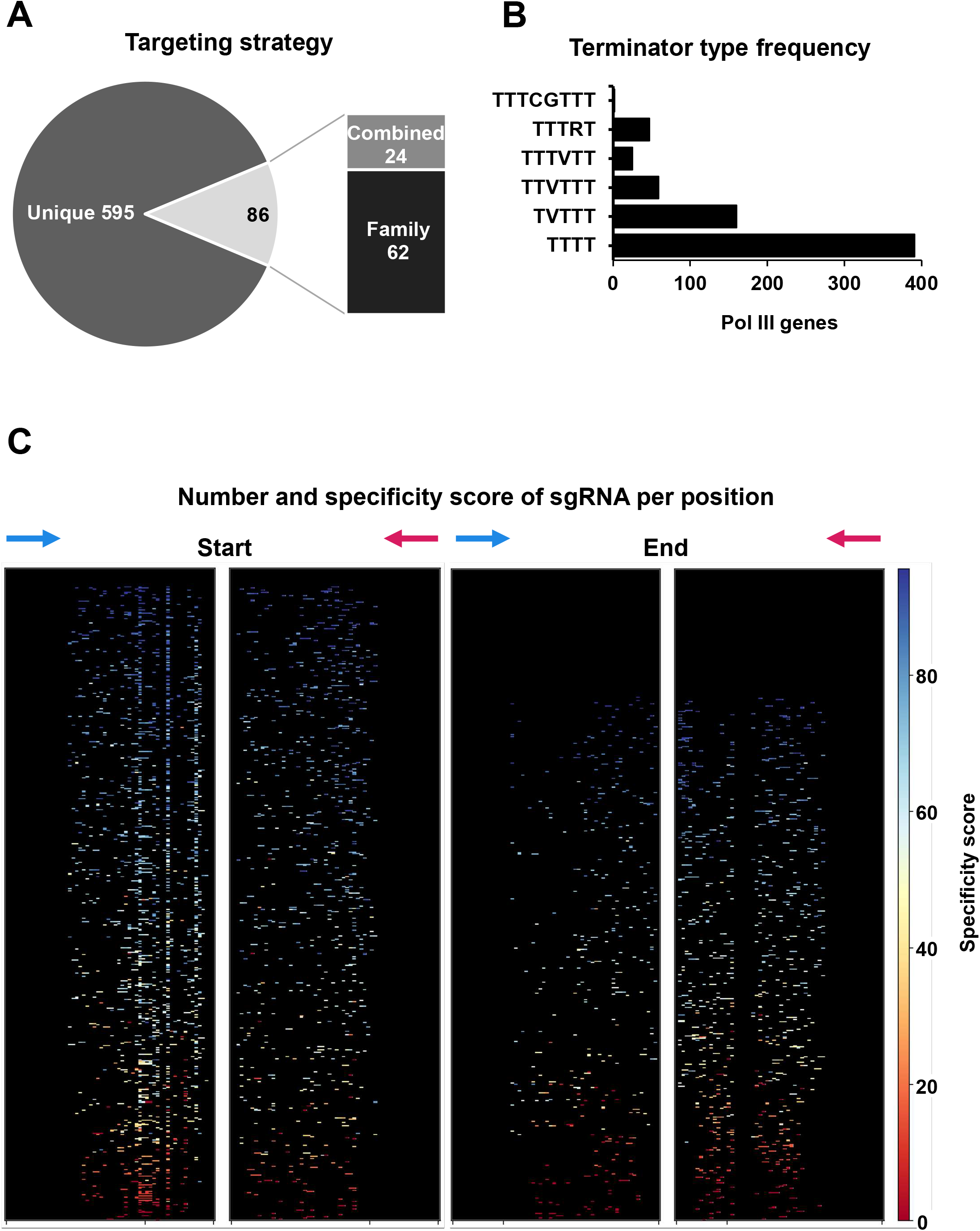
Related to Figure 2. **A**) Classification of sgRNAs according to locus specificity, including uniquely targeted loci, dual-target guides, and family-targeting guides. **B**) Frequency of canonical and non-canonical Pol III terminator motifs among loci represented in the sgRNA library. **C**) Distribution of sgRNA specificity scores across the 5′-proximal and terminator-associated targeting windows shown in Figure 2B.

**Figure S3.**
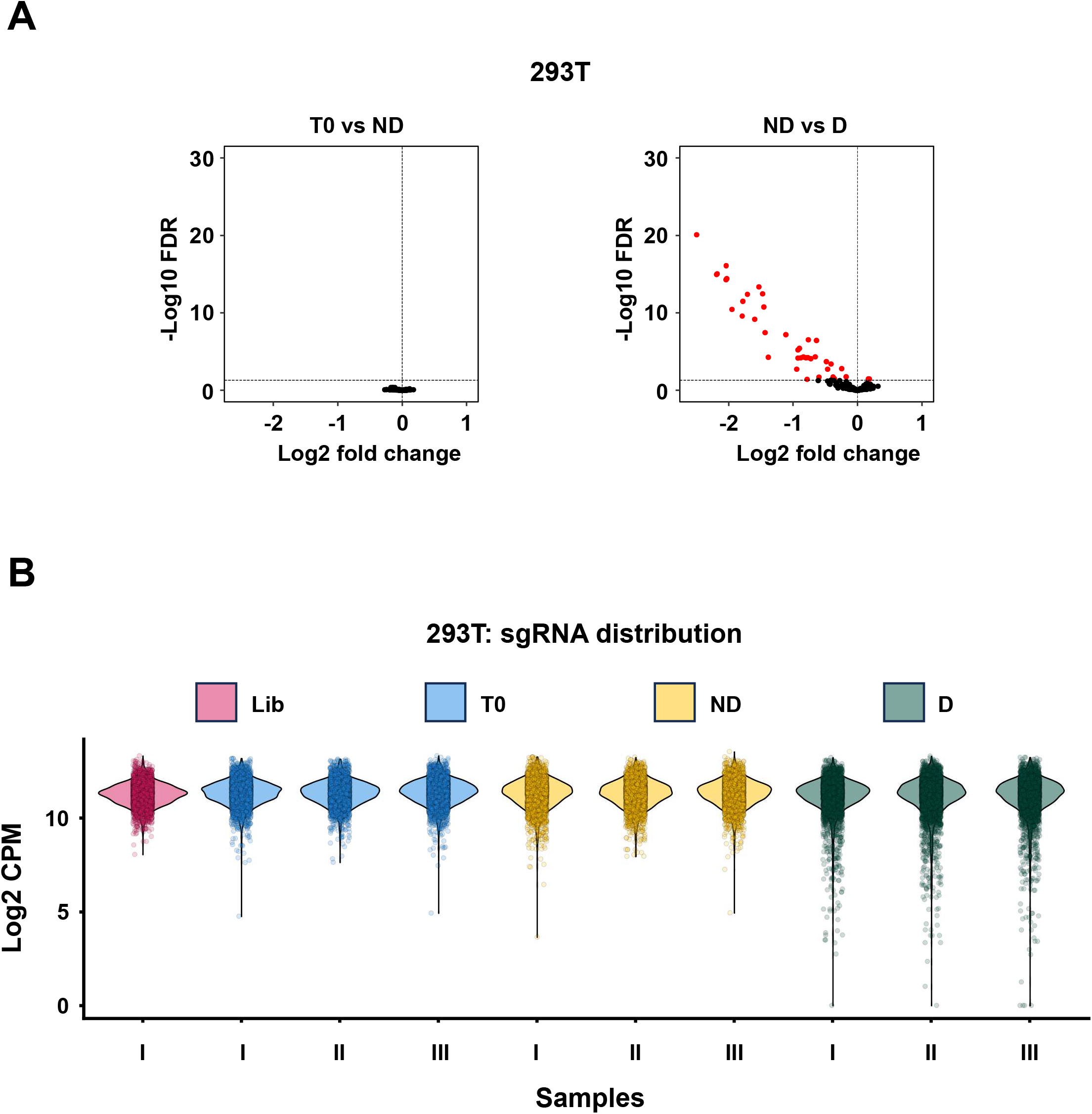
Related to Figure 3. **A**) Benchmarking volcano plots of the HEK293T screen. Left: comparison of the two control conditions, no dox (ND) versus time point zero (T0), showing no significant hits. Right: the analysis in Figure 3C recomputed against the ND control instead of T0, yielding comparable results. Significance thresholds: FDR < 0.05. **B**) Distribution of normalized sgRNA read counts across T0, untreated (ND), and doxycycline-treated (D).

**Figure S4.**
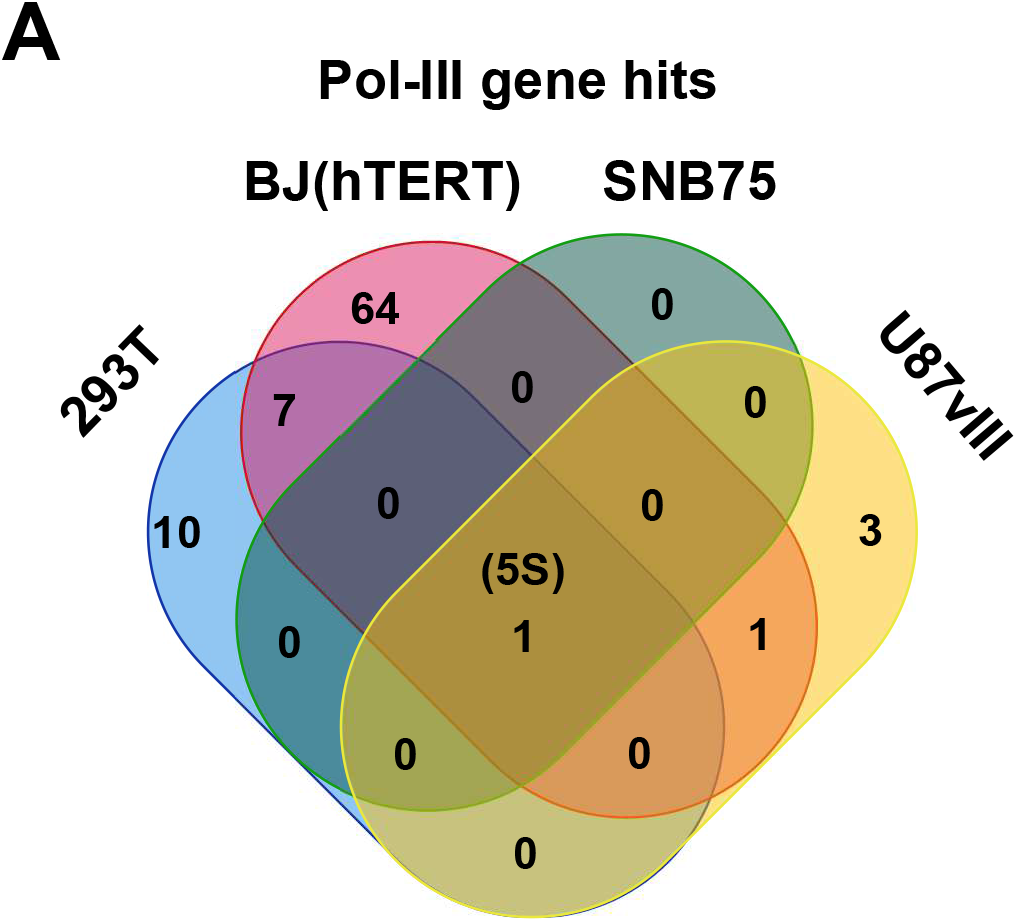
Related to Figure 4. **A**) Venn diagram showing the overlap of significant Pol III-transcribed gene hits identified in HEK293T, BJ-hTERT, SNB-75, and U87vIII cells. Hits include depleted and enriched genes with FDR < 0.05.

**Figure S5.**
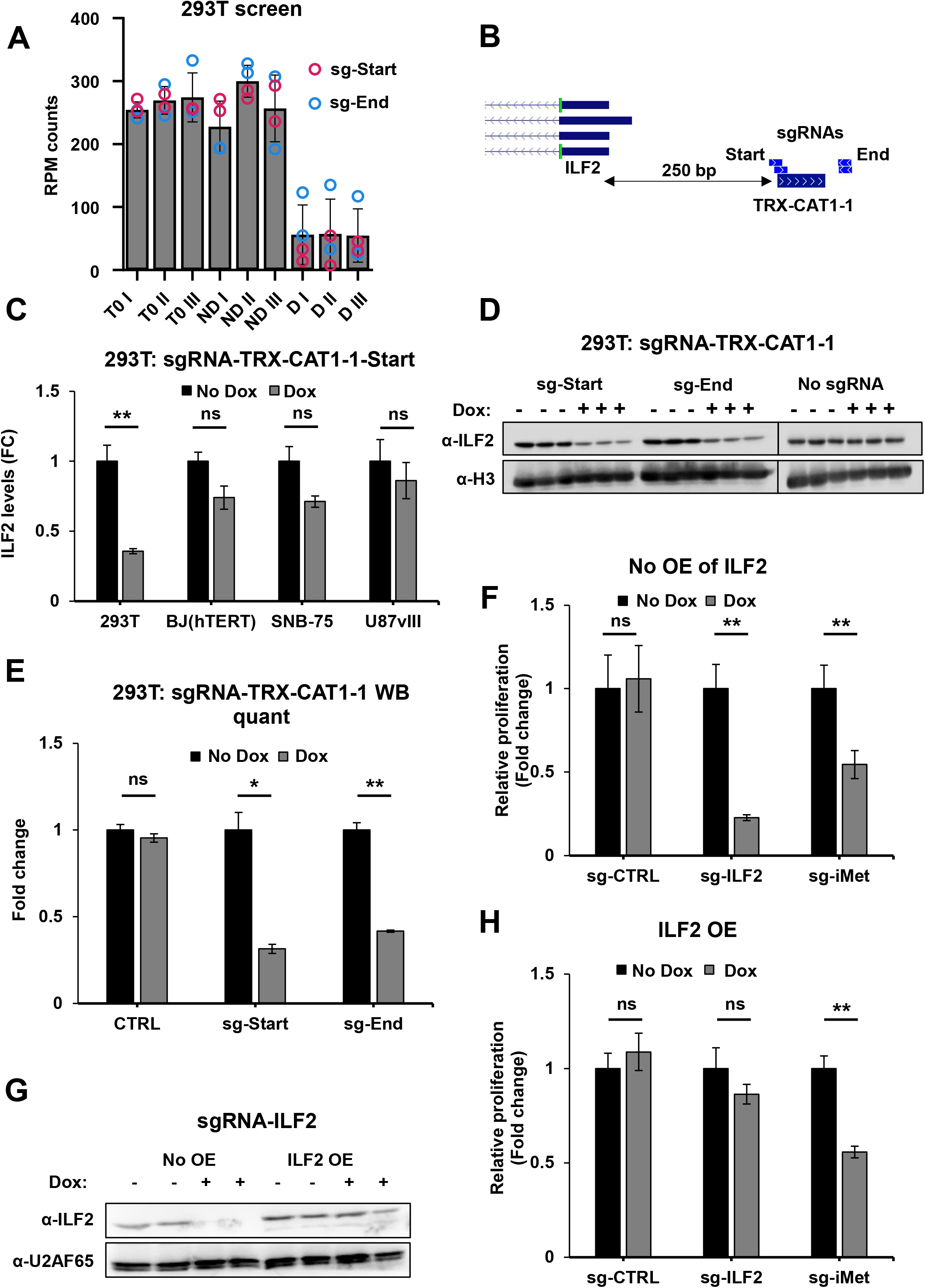
Related to Figure 5. **A**) Normalized abundance of individual sgRNAs targeting the 5′-proximal (“Start”) and terminator-associated (“End”) regions of *TRX-CAT1-1* in the HEK293T screen (n = 3). **B**) A snapshot from the UCSC Genome Browser of the *TRX-CAT1-1* and *ILF2* loci, showing their genomic proximity and the sgRNA target sites. **C**) RT-qPCR analysis of ILF2 mRNA following 7 days of *TRX-CAT1-1* repression in the indicated cell lines, normalized to PPIA mRNA levels. **D**) Immunoblot of ILF2 after one week of *TRX-CAT1-1* repression at the start and end loci. Histone H3, loading control. Representative of three independent experiments. **E**) Quantification of the blots in **D**). **F**) Relative proliferation assays of HEK293T cells after one week of *TRX-CAT1-1* or *ILF2* knockdown. Control cells were dox-treated but lacked an sgRNA (n = 5). **G**) Immunoblot confirming ectopic ILF2 variant 1 (ILF2V1; NM_004515) expression during the rescue experiment. U2AF65 was used as a loading reference. **H**) Relative proliferation assays as in **F**), with concurrent ILF2V1 overexpression (n=5). For quantitative analyses, data represent mean ± SD from independent biological replicates. Statistical significance was assessed using two-sided Welch’s t-test (*p < 0.05; **p < 0.01; ***p < 0.001).

**Figure S6.**
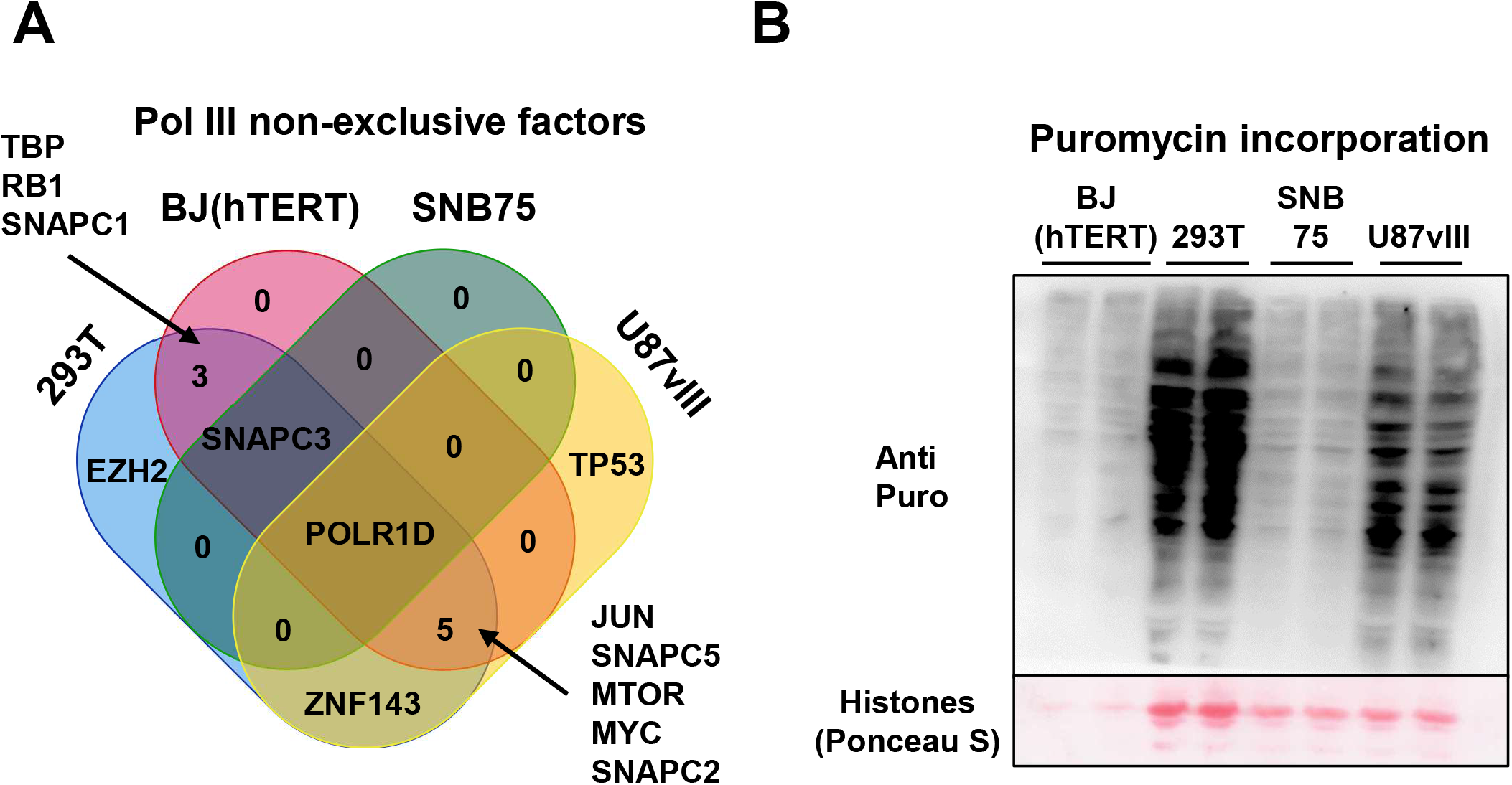
Related to Figure 6. **A**) Pol III-associated transcriptional regulators with broader functions beyond the Pol III system identified as significant hits across the tested cell lines. **B**) Puromycin incorporation assay comparing global protein synthesis across the indicated cell lines. Puromycin signal was normalized to the histone region of the Ponceau S-stained membrane as a proxy for cell number/DNA content. Quantification represents mean ± SD (n = 2 biological replicates).

### Supplementary tables

**Table S1.** List of all PCR/qPCR primers, northern blot probes, sequencing primers, and sgRNA oligo sequences.

**Table S2.** Genome-wide Pol III sgRNA library.

**Table S3:** sgRNAs normalized counts (CPM) of the sgRNAs for all CRISPRi screens.

**Table S4.** Gene-level CRISPRi screen results in HEK293T

**Table S5.** Gene-level CRISPRi screen results in BJ-hTERT

**Table S6.** Gene-level CRISPRi screen results in SNB-75

**Table S7.** Gene-level CRISPRi screen results in U87vIII

**Table S8.** List of all sgRNAs targeting multiple genes (families).

## Resource availability

NGS data from the CRISPR screen were deposited at Gene Expression Omnibus (GEO) under accession number GSE342190. The tRNA sequencing data is available at GEO with accession number GSE186736. Any additional information required to reanalyze the data reported in this paper is available from the lead contact upon request.

## Acknowledgments

We thank Keiichi Ito, Evelina Tutucci and Min Gao for their comments on the manuscript. This work was supported by a NWO Talent Programme Vidi grant [VI.Vidi.193.107] and an Amsterdam UMC Starter Grant.

## Author contributions

B.S. designed and performed experiments, analyzed the data and wrote the manuscript. Z.V. and Y.F. designed and performed experiments. A.G. designed the project, acquired funding, supervised the work and wrote the manuscript. All authors read the manuscript.

## Materials and methods

### Cell culture

HEK293T (RRID:CVCL_0063), BJ-hTERT (RRID:CVCL_6573), SNB-75 (RRID:CVCL_1706), and U-87MG-EGFRvIII^47^ cell lines were cultured at 37°C in a humidified atmosphere containing 5% CO2 in high-glucose Dulbecco’s Modified Eagle’s medium (DMEM; Sigma Life Science; D5796) supplemented with 10% fetal bovine serum (VWR; S1400-500), 1% penicillin/streptomycin (Corning; 30-002-CI). Cell lines were routinely tested and confirmed negative for mycoplasma contamination. To generate inducible CRISPRi cell lines, HEK293T, SNB-75, and U87-MG EGFRvIII cells were co-transfected with PB-TRE-dCas9-KRAB-MeCP2 vector^48^ (Addgene; #122267) and the Super PiggyBac transposase plasmid with Polyethylenimine (PEI; Polysciences; 23966-100) using a 1:2 DNA:PEI ratio. Stable integrants were selected with 300 μg/ml Hygromycin (Roche Diagnostics GmbH; 10843555001). BJ-hTERT cells were transduced with pLX-TRE-dCas9-KRAB-MeCP2-BSD^48^ (Addgene; #140690) lentivirus and selected using 2 μg/ml Blasticidin (Merck; SBR00022-1ML).

Clonal CRISPRi cell lines were generated by limiting dilution cloning. Individual clones were screened for dCas9-KRAB-MeCP2 expression by immunoblotting (**Figure S1B**), and a single clone displaying robust dCas9-KRAB-MeCP2 expression and CRISPRi activity was selected for all subsequent experiments.

sgRNAs used in this study (**Table S1**) were cloned into pL-U6-sgRNA-SFFV-Puro-P2A-EGFP^49^ (Addgene; #175037) or pKLV-U6gRNA(BbsI)-PGKpuro2ABFP^50^ (Addgene; #50946) lentiviral vectors. Stable sgRNA-expressing cell populations were generated by lentiviral transduction followed by selection with 2 μg/mL puromycin (Merck; P8833-25MG). All plasmid constructs were verified by Sanger sequencing. Cloning and sequencing primers are listed in **Table S1**.

Cell proliferation was assessed using the alamarBlue Cell Viability Reagent (Thermo Fisher Scientific; DAL1025) according to the manufacturer’s instructions. Cells were seeded in 96-well plates at 1,000 cells per well and cultured in the presence or absence of doxycycline as indicated. For *TRX-CAT1-1* repression experiments, doxycycline treatment was initiated at the time of plating and cells were cultured for 7 days before measurement. For *TRR-TCT1-1* repression experiments, cells were pre-treated with doxycycline for 2 days before plating and subsequently cultured for an additional 7 days in the presence of doxycycline. alamarBlue reagent was added directly to the culture medium, and fluorescence was measured after approximately 2 h for HEK293T, SNB-75, and U87vIII cells and after approximately 4 h for BJ-hTERT cells. Proliferation was expressed relative to the corresponding untreated control.

Global protein synthesis was assessed by puromycin incorporation. Cells at approximately 70% confluency were incubated with 5 μg/mL puromycin for 15 min. The cells were washed with 1X PBS and lysed directly in 2X Laemmli buffer without bromophenol blue (4% SDS, 20% glycerol, 10% 2-mercaptoethanol, 0.125 M Tris HCl [pH 6.8]). The lysates were boiled for 5 min at 95 °C and sonicated before immunoblot analysis.

### RNA extraction

RNA isolation was performed using TRIzol Reagent (Thermo Fischer Scientific; 15596018) as described previously^23^. Cell pellets were directly lysed in TRIzol Reagent following removal of the culture medium and washing with PBS once. After incubating for 5 minutes at room temperature, the samples were centrifuged at 12,500×g for 10 minutes at 4°C to minimize contamination from insoluble substances, including genomic DNA. Chloroform (0.2 volumes relative to TRIzol) was added to achieve phase separation, followed by thorough mixing for 30 seconds and centrifugation at 12,500 × g for 15 minutes at 4°C. The aqueous phase was collected, and RNA was isolated by adding 0.8 volumes of isopropanol and allowing it to sit at room temperature for 10 minutes, then centrifuging at 12,500 × g for 10 minutes at 4°C. The RNA pellets were washed twice with 80% ethanol, allowed to air-dry, resuspended in RNase-free water, and incubated at 55°C for 5 minutes.

### cDNA synthesis and qPCR

cDNA was synthesized from 1-3 μg of RNA template using the SuperScript IV Reverse Transcriptase (Thermo Fisher Scientific; 18090050) with random hexamers according to the manufacturer’s protocol, except for experiments assessing MIR expression levels, for which gene-specific primers were used at a final concentration of 2 μM. Briefly, the initial primer annealing and extension was performed for 10 min at RT, followed by reverse transcription for 25 min at 50°C, and enzyme inactivation for 10 min at 80°C.

Quantitative PCR (qPCR) was performed using PowerUp SYBR Green Master Mix (Thermo Fisher Scientific; A25742) in 25 µL containing 2 µL of a 1:10 dilution of the cDNA template and 0.3 μM of each primer. Amplification was performed on an Applied Biosystems 7500

Fast Real-Time PCR System. Relative transcript abundance was calculated using the 2^-ΔΔCt method and normalized to PGK1 or PPIA as indicated in the figure legends. Pre-tRNA qPCR primers were designed to anneal within 5’ leader and/or 3’ trailer sequences absent from mature tRNAs, thereby selectively detecting precursor transcripts. Primer sequences are listed in **Table S1**.

### sgRNA library design and cloning

Pol III-transcribed loci were compiled from tRNAscan-SE^10^ for tRNA genes and from RepeatMasker^51^, and UCSC annotations for other Pol III-transcribed elements. Candidate loci were restricted to transcriptionally active genes based on Pol III ChIP-seq occupancy in IMR90-hTERT fibroblasts^52^. When a single ChIP-seq peak overlapped multiple annotated Pol III loci, tRNA annotations were prioritized for locus assignment. CRISPOR^53^ and CRISPick^54,55^ databases were used to select highly specific (MiT score >= 40) sgRNAs overlapping with the Pol III-transcribed loci. Because Pol III transcription termination sites are incompletely annotated, canonical and non-canonical^58^ terminator motifs were identified computationally using the pattern recognition software COMPASSS^56^, and their genomic coordinates were used to define terminator-associated sgRNA targeting windows.

Targeting windows were defined empirically from the sgRNA tiling experiments shown in **Figure 1**. For type II and type III Pol III genes, repression efficiency was measured by RT-qPCR for sgRNAs positioned across upstream, gene-body, and downstream regions. These data were used to define the 5′-proximal and terminator-associated targeting windows used for library design (**Figure 2A**). Up to eight sgRNAs were selected per locus to balance library size with guide redundancy. Loci with fewer available guides were included when at least two sgRNAs meeting the design criteria could be identified (**Figure 2D**). Guides predicted to target multiple Pol III loci were retained when shared targeting was intentional. These were annotated either as family-targeting sgRNAs or as dual-target guides, depending on the number of matched loci (**Table S8**). Positive-control sgRNAs targeting established essential genes were selected from published CRISPRi datasets, including MCM2, GEMIN5, CENPA, INTS9, POLR1D, RPL11, RPS8, RPL30, PSMD1, COPZ1, and POLR2A.

sgRNAs targeting Pol III machinery components were selected from published CRISPRi guide resources, and non-targeting sgRNAs were drawn from a previously validated control set^19,55,57–59^. The final library contained 4110 sgRNAs that targeted genes. Thirty-four candidate Pol III loci were excluded because fewer than two sgRNAs meeting the design criteria could be identified.

The pooled sgRNA library was synthesized as single-stranded oligonucleotides with 5’-ATATCTTGTGGAAAGGACGAAACACCGG and 3’-GTTTTAGAGCTAGAAATAGCAAGTTAA flanking sequences (GenScript). Oligonucleotides were PCR-amplified using ArrayF and ArrayR primers (**Table S1**) and cloned into pL-U6-sgRNA-SFFV-Puro-P2A-EGFP (Addgene; #175037) using NEBuilder HiFi DNA Assembly Master Mix (New England Biolabs; E2621S). To maintain library complexity, two independent transformations were performed using XL10-Gold Ultracompetent Cells (Agilent; 200315). Transformation yielded approximately 2.7 × 10^5^ colonies, corresponding to ∼66-fold representation of the 4,110-sgRNA library. Colonies were pooled by scraping, and plasmid DNA was isolated by maxiprep (Thermo Fisher Scientific; K0492).

### Lentivirus production and transduction

Lentiviral particles were produced in HEK293T cells cultured in poly-L-lysine-coated plates (Merck; A-005-C). Cells were co-transfected with the lentiviral transfer plasmid, psPAX2 (Addgene #12260), and pMD2.G (Addgene #12259) plasmids at a DNA mass 2:1:1 ratio using polyethylenimine (PEI; Polysciences, #23966-100) with a 1:2 DNA:PEI ratio in OptiMEM (Thermo Fisher Scientific; 31985062). Medium was replaced 6 h after transfection, and viral supernatants were collected at 24, 48, and 72 h after medium replacement, pooled, passed through a 0.45 μm filter, and used directly for transduction.

Target cells were transduced in the presence of 8 μg/mL polybrene (Santa Cruz Biotechnology; sc-134220A). For pooled CRISPRi screens, lentiviral particles carrying the sgRNA library were produced using scaled-up transfections in 15-cm dishes. The viral input was adjusted to achieve a multiplicity of infection (MOI) of approximately 0.3-0.4, estimated from the percentage of GFP-positive cells using an IncuCyte ZOOM live-cell analysis system (Essen BioScience), to favor single-vector integration events.

### CRISPRi screening procedures

The cells were seeded at 4 x 10^6^ per sample and were transduced 24 h later with the pooled sgRNA library at an estimated MOI of ∼0.3–0.4 corresponding to approximately 30% transduction efficiency. Following puromycin selection, T0 samples were collected and populations were maintained at a minimum of 4 × 10^6^ cells per sample throughout the screen to preserve library representation. This corresponded to ∼970-fold representation of the 4,110-guide library after selection. For the initial HEK293T benchmark screen, cells were subsequently cultured in the presence or absence of 2 µg/mL doxycycline (Sigma-Aldrich; D9891). Subsequent screens were performed with doxycycline-treated populations and analyzed relative to T0. Cells were maintained for approximately 20 population doublings before collection. The genomic DNA extraction was performed from 4 x 10^6^ cells per sample. All screens were performed in three independent biological replicates.

### Genomic DNA extraction and sequencing library preparation

Cell pellets were resuspended in 400 μL of lysis buffer (50 mM Tris pH 8.0, 100 mM EDTA, 100 mM NaCl, and 1% SDS) supplemented with 2 mg of proteinase K (Thermo Fisher Scientific; #4333793) and incubated overnight at 50°C. The lysates were diluted to 1 mL with 10 mM Tris (pH 8.0) and extracted with 1 volume of Phenol:Chloroform/Tris Buffer (Thermo Fisher; 11886714). The aqueous phase was subsequently treated with RNase A (Thermo Fisher Scientific; EN0531) for 2h at 50°C prior to extraction with water-saturated chloroform. Genomic DNA was finally precipitated with sodium acetate and isopropanol. DNA pellets were washed with 70% ethanol and resuspended in 10 mM Tris-HCl (pH 8.0).

To maintain library representation during sequencing library preparation, 15 μg gDNA was used per sample. Integrated sgRNA cassettes were amplified in six parallel PCR reactions containing 2.5 μg gDNA each using Q5 Hot Start High-Fidelity DNA Polymerase (New England Biolabs; M0494S). The first PCR was performed for 21 cycles to amplify sgRNA-containing fragments and introduced partial sequencing adapters. Parallel reactions were pooled, and one-fifth of the reaction was analyzed by 8% polyacrylamide gel electrophoresis to verify amplification. A second PCR of 6 cycles was used to incorporate Illumina adapters and sample-specific barcodes. Final sequencing libraries were purified using the GeneJET Gel Extraction and DNA Cleanup Micro Kit (Thermo Fisher Scientific; K0832), eluted in 10 mM Tris-HCl (pH 8.0), quality controlled by 8% polyacrylamide gel electrophoresis, and sequenced on an Illumina NovaSeq 6000 platform. Primer sequences are listed in **Table S1**.

### CRISPRi screen data analysis

Raw sequencing reads (FASTQ) were demultiplexed and mapped to the sgRNA reference library using Guide Counter^60^, allowing no mismatches between sequencing reads and sgRNA reference sequences. The resulting sgRNA count matrix was used for all downstream analyses. Sequencing quality was assessed based on total read counts, mapping efficiency, and sgRNA representation across samples.

Gene-level effects were estimated using the Maximum Likelihood Estimation (MLE) module implemented in MAGeCK^59,61^. For each screen, sgRNA count tables from doxycycline-treated samples were compared with the corresponding reference population using a design matrix that modeled treatment effects across biological replicates. In the HEK293T benchmark screen, analyses performed using either untreated (ND) or initial (T0) reference samples produced highly concordant results (**Figure S3A**); therefore, T0 was used as the primary reference condition for subsequent screens.

To account for differences in sequencing depth and sample representation, MAGeCK normalization was performed using non-targeting control sgRNAs with the ––control-sgrna parameter. Gene-level effects were summarized as beta scores, with negative values indicating depletion and positive values indicating enrichment relative to the reference population. Statistical significance was assessed using the Wald test implemented in MAGeCK, and false discovery rates (FDR) were calculated using the Benjamini-Hochberg procedure. Genes with an FDR < 0.05 were considered significant. Published tRNA abundance data from Scheepbouwer et al.^17^ were used for comparison with screening results.

### Extract preparation and Western blotting

For whole-cell lysates, cells were lysed in RIPA buffer (150 mM NaCl, 1% Nonidet P-40, 0.5% DOC, 0.1% SDS, and 50 mM Tris pH 7.4) supplemented with a protease inhibitor cocktail (Bioke; 5871). Protein concentration was determined using the Pierce BCA protein assay kit (Thermo Fisher Scientific; 23225). Equal amounts of protein were mixed with Laemmli sample buffer supplemented with β-mercaptoethanol and denatured at 90°C for 5 min.

For cellular fractionation, cells were pelleted at 500 × g for 3 min at room temperature and washed twice with ice-cold PBS. Pellets were resuspended in 250 µL of a 1:1 mixture of 2× NUN buffer (50 mM HEPES-NaOH, pH 7.6; 0.6 M NaCl; 2% NP-40; 2 M urea; 2 mM DTT) and nuclear lysis buffer (10 mM HEPES-KOH, pH 7.6; 100 mM KCl; 0.1 mM EDTA; 0.15 mM spermine; 0.5 mM spermidine; 10% glycerol; 0.5 mM DTT). Samples were incubated on ice for 20 min and centrifuged at 17,000 × g for 20 min at 4°C. The supernatant, containing the soluble cytoplasmic and nucleoplasmic fraction, was collected. The remaining pellet was washed twice with the same NUN/Nuclear lysis buffer mixture and centrifuged at 10,000 × g for 5 min. The chromatin-associated fraction was extracted from the pellet in urea buffer (8 M urea; 200 mM Tris-HCl, pH 6.8; 1 mM EDTA; 5% SDS; 1.5% DTT) at one-tenth the volume used for the soluble fraction and sonicated until fully solubilized.

Proteins were separated by SDS-PAGE and transferred onto Immobilon-P PVDF Membranes (Merck; IPVH00010). Membranes were blocked in 5% skim milk in TBST for 1 h at room temperature and incubated overnight at 4°C with primary antibodies. Following washing, membranes were incubated for 1 h at room temperature with HRP-conjugated secondary antibodies (Cell Signaling Technology; 7074S and 7076S). Protein bands were detected using SuperSignal West Pico Plus chemiluminescent substrate (Thermo Fisher Scientific; 34580) and imaged using a Uvitec chemiluminescence system. Primary antibodies and dilutions are listed in **Table S1**.

### Northern blotting

Northern blotting was performed as previously described ^17,23^. Total RNA was denatured for 1 min at 90°C in RNA loading dye (New England Biolabs; B0363S) and separated on a 10% urea-polyacrylamide gel in 1X TBE buffer. RNA loading was assessed by staining gels with SYBR Gold nucleic acid gel stain (Thermo Fisher Scientific; S11494) for 20 minutes at RT.

RNA was transferred onto a positively charged nylon membrane (CYTIVA; RPN203B) in 0.5X TBE buffer and UV-crosslinked using a Stratalinker 2400 (Stratagene; 15933). The membrane was pre-hybridized with ULTRAhyb-Oligo buffer (Thermo Fisher Scientific; AM8663) at 42°C for 1 hour and hybridized overnight with 50 nM biotinylated probes at 37°C. Following hybridization, the membrane was washed twice in low-stringency wash buffer (1X SSC, 0.5% SDS) and blocked in 2X SSC, 0.5% SDS, 3% BSA for 15 min. Biotinylated probes were detected by incubation with HRP-conjugated streptavidin (1:40,000; Thermo Fisher Scientific, N100) for 30 min. The membrane was subsequently washed once with ABS buffer (2X SSC, 10% BSA, 1% TritonX-100) for 5 min and rinsed twice with 2X SSC prior to signal detection using SuperSignal West Pico Plus chemiluminescent substrate (Thermo Fisher Scientific, 34580) and imaged on a Uvitec chemiluminescence system. Band intensities were quantified using ImageJ. All Northern blot probe sequences are listed in **Table S1**.

### ILF2 overexpression and rescue assay

ILF2 variant 1 cDNA (NM_004515) was generated by reverse transcription using a transcript-specific primer (**Table S1**). The cDNA was first cloned downstream of a CMV promoter in pEGFP-C1. The CMV-ILF2 cassette was subsequently amplified by PCR and subcloned into pCWX-PGK-BSD^62^ (Addgene; #114316). The resulting ILF2 overexpression construct was introduced into HEK293T cells by lentiviral transduction, and ILF2 expression was verified by immunoblotting.

For rescue experiments, HEK293T CRISPRi cells with or without ectopic ILF2 expression were transduced with sgRNAs targeting either ILF2 or the 5′-proximal region of TRX-CAT1-1. CRISPRi was induced with doxycycline, and proliferation was measured using alamarBlue in five independent replicates per condition. Growth of ILF2-overexpressing cells was compared with the corresponding non-overexpressing controls.

### Statistical analysis

Statistical analyses for validation experiments were performed using independent biological replicates, with exact replicate numbers indicated in the figure legends. Band intensities were quantified using FIJI/ImageJ^63^ (version 1.54r). For qPCR, immunoblot quantification, Northern blot quantification, and proliferation assays, statistical significance was assessed using two-sided Welch’s t-tests unless otherwise indicated. Data are presented as mean ± SD. Statistical analysis of pooled CRISPRi screens was performed as described above using MAGeCK, with FDR < 0.05 considered significant.

